# Systematic mapping of chromatin dysregulation driven by viral transcriptional regulators at scale

**DOI:** 10.64898/2026.05.17.725756

**Authors:** James E. Corban, Keelin Reilly, Jaice T. Rottenberg, Andrea Hunger, Max Frenkel, Juan Fuxman-Bass, Srivatsan Raman

## Abstract

Viral transcriptional regulators (vTRs) reprogram host cell transcriptional and epigenetic networks to promote infection, persistence, and oncogenic transformation. Despite their broad essentiality to viral pathogenesis, thousands of putative vTRs remain functionally uncharacterized. Here we apply PROD-ATAC, a pooled single-cell epigenomic screening platform, to map chromatin accessibility dysregulation induced by over 100 vTRs in a single assay. Our screen identified dozens of vTRs which directly and indirectly remodel chromatin, including variants with no previously reported epigenetic activity. Comparative analyses revealed that unrelated vTRs frequently opened chromatin regions associated with the same host transcription factor networks, including the NF-κB, AP-1, p53, and Sp1/KLF families. Integration of chromatin accessibility and transcriptomic datasets highlighted that epigenetic perturbations often coincide with downstream gene expression changes. By adapting PROD-ATAC to incorporate small-molecule perturbations, we demonstrate that targeted inhibition of host features (including the MAPK signaling cascade) can illuminate host dependencies for vTR-induced epigenomic dysregulation. These results establish a scalable framework for discovering virus-host chromatin interactions and suggest that epigenetic manipulation of host regulatory networks is a widespread and conserved function of many diverse vTRs.

## Introduction

Viruses are the most abundant and evolutionarily dynamic biological entities on Earth^1,2^. Cells across all domains of life are susceptible to infection by diverse viral species which drive pathogenesis through distinct strategies^3,4^. A defining feature of many viral infection cycles is their capacity to reprogram host cellular environments to promote viral replication, persistence, and transmission. Central to this process are viral transcriptional regulators (vTRs), a diverse category of viral proteins that directly or indirectly modulate host transcriptional networks^5^. By targeting key host transcription factors (TFs), chromatin remodeling complexes, and signaling pathways, vTRs can dramatically reshape cellular regulatory landscapes^5–7^. These activities contribute to aspects of viral pathogenesis, such as immune system stimulation, cell stress, and oncogenic transformation^8^. Several well-characterized vTRs, including multiple oncoproteins expressed by the human tumor viruses Epstein-Barr virus (EBV)^6^, Kaposi’s sarcoma-associated herpesvirus (KSHV)^9^, and human T-cell leukemia virus (HTLV)^10^, drive pathogenesis by remodeling epigenetic landscapes.

Despite their biological and clinical significance, the epigenetic roles of most vTRs remain poorly understood. The entire viral proteome space encodes thousands of putative transcriptional regulators spanning diverse viral families and genome architectures^5,8,11,12^. However, functional characterization of these proteins has historically relied on low-throughput, arrayed experimental approaches such as chromatin immunoprecipitation sequencing (ChIP-seq), bulk ATAC-seq, or targeted transcriptional assays^13^. While these studies are essential and have yielded valuable insights into vTR-driven epigenetic dysregulation strategies, these targeted studies do not approach the scale of viral epigenetic regulators that remain understudied. Furthermore, accurate epigenetic and transcriptomic comparisons across experiments are frequently confounded by sample-to-sample variations and batch effects^14,15^. As a result, our current understanding of how viral proteins reshape host chromatin landscapes remains limited and biased toward a relatively small quantity of extensively studied vTRs^5^.

We sought to establish a large-scale survey of vTR-induced chromatin dysregulation to systematically characterize the diverse mechanisms by which these abundant viral proteins hijack host regulatory landscapes. Recent advances in single-cell functional genomics have enabled the high-throughput characterization of gene-based perturbations at unprecedented scales^16,17^. Particularly relevant are the Perturb-seq^18^ and Spear-ATAC platforms^19^, pooled variant single-cell CRISPR-based screening platforms coupled to transcriptomic and epigenomic readouts, respectively. However, these screening frameworks only enable the assignment of short gRNA sequences to specific cell identities and cannot recover the genotypes of vTRs or other protein-coding length variants. Extensions of the Perturb-seq framework (sc-eVIP) have enabled pooled protein-coding perturbation analyses^20^, yet analogous strategies to recover variant identity alongside epigenomic-based readouts are lacking. Lentiviral delivery of protein-coding libraries offers an alternative strategy for pooled variant screening of epigenetic dysregulation, yet complications can arise through semi-random genomic integration and template switching during lentiviral packaging. Resultant barcode-variant assignment errors and cell-to-cell variability can obscure genotype-phenotype relationships in single-cell assays^21,22^.

We addressed these challenges by applying PROD-ATAC, our recently developed method which maps the effects of protein-coding variants to their resultant chromatin perturbations via single-cell ATAC-seq^23^. PROD-ATAC employs a recombinase-based integration system that inserts protein variants alongside DNA barcodes into a single defined genomic landing site, ensuring that each cell expresses a single variant in an identical cell context. Following inducible expression of the variant library, chromatin accessibility landscapes are measured using a modified single-cell ATAC-seq workflow that simultaneously captures variant-identifying barcodes^23^. This strategy links vTR genotypes to single-cell chromatin accessibility profiles, facilitating large-scale mapping of epigenetic phenotypes across diverse vTRs in a shared cell background.

Using the PROD-ATAC platform, we performed a pooled epigenomic survey of more than 100 vTRs derived from 15 viral families with distinct evolutionary backgrounds. This assay used only ∼140,000 cells (approximately the quantity needed for a single bulk ATAC assay) to generate chromatin accessibility landscapes for over 100 variants. The resulting dataset revealed that chromatin remodeling is a common and often heterogeneous property of vTRs. While several well-studied vTRs produced expected chromatin dysregulation, many other variants with no reported chromatin-modifying activity emerged as potent epigenetic modulators^5^. Additionally, we highlight examples of vTR homologs which exhibited vastly different chromatin-remodeling strengths. Comparisons of accessibility landscapes further revealed similar functional relationships between vTRs from unrelated viral lineages. Several structurally and phylogenetically unrelated vTRs generated similar chromatin accessibility shifts and opened chromatin enriched for binding sites of key host TF families, including NF-κB, AP-1, p53, and Sp1/KLF. Pathways regulated by these TF families control fundamental cell processes including immune signaling, stress responses, and proliferative control^24,25^. These results suggest that vTRs repeatedly converge on shared host pathways to reprogram cellular states. Integrating our epigenomic datasets with bulk transcriptomic analyses revealed that chromatin perturbations often co-occurred with substantial gene expression shifts, reinforcing the functional significance of the measured chromatin accessibility changes.

In this work, we separately extended the PROD-ATAC framework to incorporate small-molecule perturbations. This perturbation assay constituted the first reported pooled screening of protein-coding variants in the presence of targeted inhibitors with a chromatin accessibility readout. Screening vTRs alongside MAPK- and BET-inhibiting molecules revealed drug-dependent shifts in chromatin accessibility landscapes^26,27^. Among other findings, these experiments uncovered signaling dependencies that distinguish direct chromatin-pioneering vTRs from those reliant upon host signaling cascades. The PROD-ATAC drug perturbation screens demonstrated that pooled variant epigenomic screening can systematically resolve host-vTR regulatory interactions.

Together, these datasets establish a scalable framework for systematically interrogating the epigenetic activities of viral proteins. PROD-ATAC overcomes key limitations of traditional arrayed approaches by enabling high-throughput mapping of chromatin accessibility landscapes induced by diverse vTRs within a unified cellular context. Our systematic characterization of vTR-driven chromatin modification provides critical insights into the mechanisms by which viruses reprogram host cells and lays the groundwork for future atlas-scale surveys of the epigenomic perturbations induced by viral proteins.

## Results

### Pooled characterization of epigenetic disruptions induced by vTRs in a shared cellular context at scale

The measurement of vTR-driven chromatin dysregulation effects required a platform suitable to the scale and diversity of the thousands of identified vTR proteins^5,7^. Multiple existing methods facilitate the high-throughput characterization of epigenomic or transcriptomic shifts induced by CRISPR-based perturbations, yet these strategies are restricted by their size (guide RNAs) and scope (contexts in which target gene(s) are expressed)^18,19^. Lentiviral-delivered sequences integrate semi-randomly into target genomes, introducing cell-to-cell variability which (especially in single-cell assay contexts) confounds the fidelity of pooled variant screening^21,22^. Additionally, template switching during lentiviral production contributes to variant barcode shuffling, a problem exacerbated with sequences several kilobases long (which includes many vTRs)^28^. The PROD-ATAC method satisfies each of these concerns by employing a Bxb1 recombinase-based system which inserts barcoded single protein-coding variants of arbitrary size into clonal 293T acceptor cells (293T-LP) that each encode a single pre-integrated landing site^23^ (Fig 1a). Once the protein-coding variants of choice are recombined into the 293T-LP cells, they undergo selection to yield a pooled population of cells that inducibly control the expression of the barcoded variants. The successfully recombined landing sites include a set of twin DNA barcodes in the 3’ untranslated region (UTR) that uniquely identify each variant. The pooled cell library is subsequently screened using an adapted 10x Genomics EpiATAC protocol which synthesizes the 3’ UTR barcodes alongside the 10x-provided nuclei barcodes. This twin barcoding scheme enables the linkage between variant genotypes detected in each cell and their corresponding epigenetic phenotypes as measured by scATAC-seq^23^.

**Figure 1:**
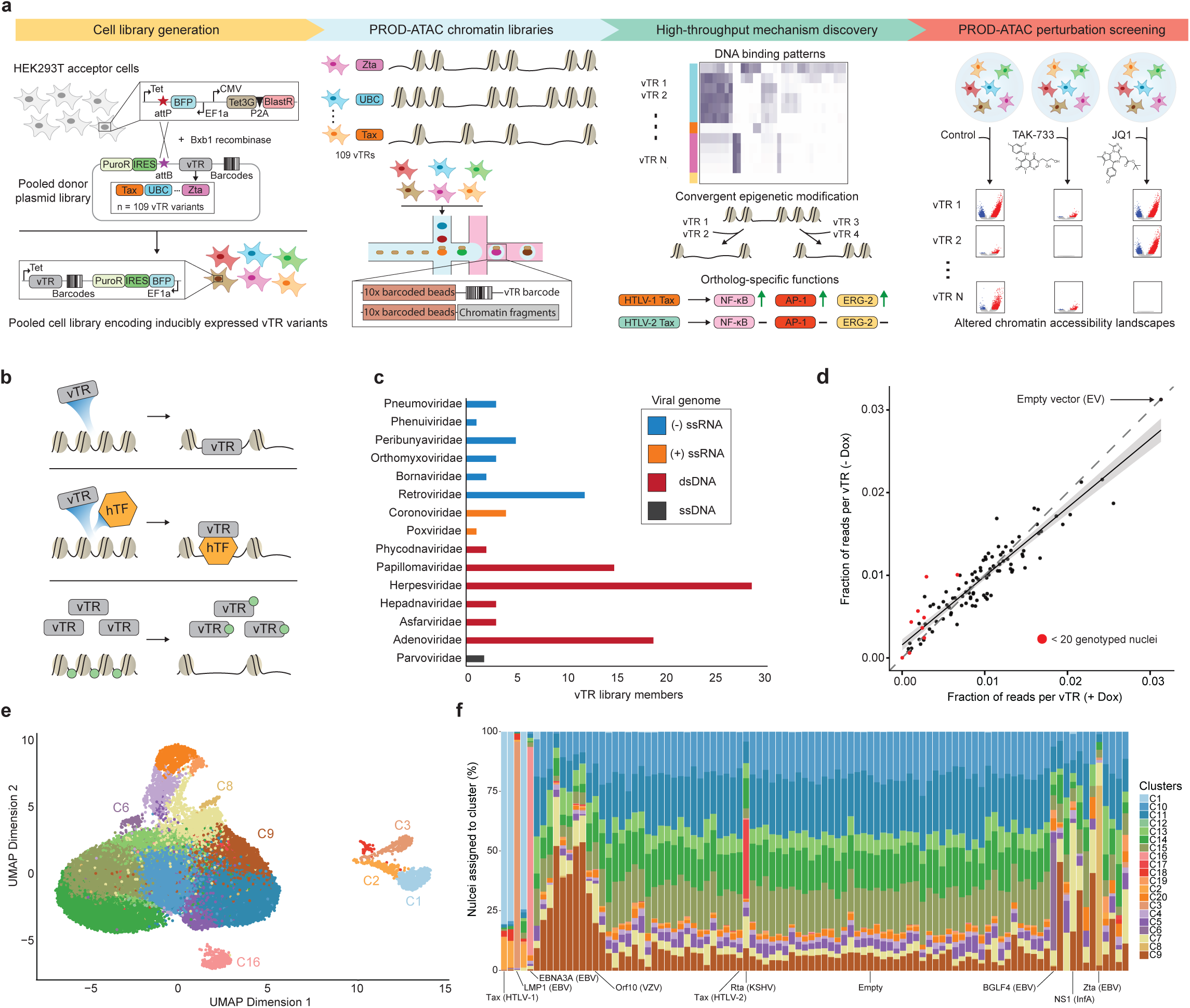
PROD-ATAC facilitates high-throughput characterization of epigenetic perturbations. **(a)** Overview of the single-cell PROD-ATAC workflow. A donor plasmid library encoding a set of 110 barcoded vTR variants (including an EV control) was recombined into 293T-LP-B4 landing pad cells. Variant expression was induced for 96 hrs, and the resultant chromatin dysregulation patterns alongside corresponding vTR identities were resolved for individual nuclei. Subsequent PROD-ATAC screening of a condensed vTR library revealed unique drug-by-variant chromatin dysregulation phenotypes. **(b)** Representations of direct (DNA-binding) and indirect (hTF interactions, host cell protein sequestration) mechanisms by which vTRs alter host cell chromatin accessibility states. **(c)** Viral families represented in the 110 screened variants, colored by viral genome nucleic acid content. **(d)** Distribution of sequencing read proportions for 110-member library after 96 hours of growth with and without doxycycline induction, with selected variants and EV control labeled. Linear regression and 95% confidence interval are shown. **(e)** UMAP embedding post latent semantic indexing (LSI) for 40,835 genotyped variant-expressing nuclei. Filtered to include only vTRs with >20 assigned nuclei (n = 100 variants). **(f)** Percentage of variant nuclei assigned to each of the 20 identified clusters displayed in (Fig 1e).

We curated a set of 110 vTRs to deeply characterize via PROD-ATAC which included proteins indicated by previous literature and homology analyses to be direct chromatin interactors and/or significant transcriptional effectors^5^ (Supplementary Table 1). Some of these vTRs are known to drive epigenetic dysregulation by directly binding host DNA, such as Zta (EBV)^29^. Many other variants indirectly alter host chromatin through interactions with host TFs and cofactors, inhibiting chromatin architectural proteins, or by other currently undefined mechanisms (Fig 1b)^5,7^. The library members represented 15 distinct viral families with diverse phylogenetic backgrounds and genome structures (Fig 1c). Most vTRs were sourced from dsDNA eukaryotic viruses. Herpesviruses, papillomaviruses, and adenoviruses (28, 20, and 18 variants, respectively) together comprised 60% of the total vTR library (Fig 1c). The preferential inclusion of dsDNA virus-derived vTRs was due to the comparative abundance of confirmed direct/indirect dsDNA interacting proteins relative to those in viruses with non-dsDNA genome architectures^5^. The 110-member vTR library was cloned into the plasmid donor constructs and assembled into our 293T-LP cells for PROD-ATAC screening as previously described^23^. A pooled donor plasmid library encoding the 110 variants was transfected into clonal 293T-LP cells. The vTR sequences on the donor plasmids were recombined via Bxb1 into the 293T-LP landing sites, and cells were selected to generate a pooled 293T-LP population which inducibly expressed a single vTR variant per cell. A high-throughput proliferation screen demonstrated that vTR library expression in 293T-LP cells introduced relatively minor proliferative defects for only a select few variants, with particularly strong defects noted for several peribunyavirus non-structural proteins (Fig 1d). As the average overall proliferative shifts were minute, we determined that library expression would not substantially obscure variant representation during downstream analyses.

PROD-ATAC sequencing of the vTR library yielded 137,491 high-quality nuclei and a dial-out sequencing dataset which linked variant genotypes to the barcoded ATAC libraries unique to each cell. Stringent genotyping confidence cutoffs assigned vTR variant genotypes to 43,798 nuclei, which equated to 31.8% of the total captured cells (Supplementary Table 2). This value is comparable to previous Spear-ATAC and PROD-ATAC efficiencies and reflects the difficulty of detecting single-copy molecular identifiers relative to analogous RNA-seq genotyping efforts, in which gene expression amplifies barcodes or gRNAs by orders of magnitude prior to sequencing. The assigned nuclei represented all 110 members within the input vTR library. An average of 398 genotyped nuclei (405 median) were assigned per variant. Key quality control metrics, including fragments within chromatin peaks and transcription start site (TSS) enrichment scores, were similar across the variant library (Supplementary Fig 1).

The initial qualitative assessment of vTR impact on cell chromatin accessibility included mapping the distribution of genotyped nuclei across dimension-reduced space. While any dimensionality reduction strategy can obscure the true space between cells, a two-dimensional projection of massive chromatin accessibility datasets (such as those produced via PROD-ATAC) can identify high-level relationships between dramatic cell phenotype changes which occur due to vTR expression. We performed iterative latent semantic indexing (LSI) coupled with uniform manifold approximation and projection (UMAP) reduction analyses on all 137,491 high-quality nuclei^30^. Subsequent unsupervised graph-based clustering of the dimension-reduced chromatin accessibility data with Seurat^31^ generated 20 distinct clusters (Fig 1e). For all downstream analyses, we filtered the vTR library members to include the 100 total variants (99 vTRs and empty vector control) for which we recovered more than 20 genotyped nuclei. We measured each variant’s distribution across the 20 identified clusters to identify the vTRs which dramatically deviated from control cell states. Empty vector (EV) cells occupied several clusters, which represented the natural cell state heterogeneity captured by single-cell assays^32^ (Supplementary Fig 2). By comparing the percent cluster occupancy of each vTR to that observed for the EV cells, we noted multiple variants which produced substantial chromatin landscape shifts. For example, 75.3% of Zta (EBV) nuclei occupied cluster C8, in which no other variant exceeded 1% occupancy (Fig 1f). Additionally, between 74.4-80.5% of nuclei assigned to three Tax (HTLV-1/3/4) homologs presided in C1, and 91.0% of EBNA3A (EBV) nuclei were present in C16 (Fig 1f). No EV control cells occupied these three clusters. These dimension reduction analyses demonstrate that PROD-ATAC can resolve broad-stroke chromatin dysregulation shifts induced by vTRs and initiates the process of classifying variants by patterns of epigenetic dysregulation.

### vTRs alter cell chromatin accessibility landscapes

We quantified the chromatin-modulating capabilities of all vTR library members by comparing the chromatin accessibility profiles for each variant to the EV control. First, chromatin peaks were called via MACS3^33^ for cells grouped by vTR genotypes. We then produced pseudobulk replicates for each variant. Pairwise differential analysis between each of the 99 filtered vTRs and the EV control identified peaks which were significantly dysregulated due to vTR activity (Fig 2a and Supplementary Fig 3). Our large-scale PROD-ATAC screen identified 33 vTRs which significantly dysregulate host cell chromatin accessibility. These disruptions ranged from dramatic genome-wide remodeling (Tax (HTLV-1) = 40,677 significant peaks) to subtle variations (L2 (HAdvF) = 30 significant peaks). Chromatin-modifying vTRs included members from seven distinct viral families, including dsDNA (Herpesviridae, Papillomaviridae, Adenoviridae, Asfarviridae, and Phycodnaviridae) and ssRNA (Orthomyxoviridae and Retroviridae) viruses^1,5^ (Fig 2a). Several vTRs previously reported to pioneer cell chromatin, including Zta (EBV)^29^ and Rta (KSHV)^34^, produced hundreds to thousands of increased accessibility peaks. Many other vTRs exhibited chromatin accessibility changing activities which had not previously been reported. These novel variants included BGLF4 (EBV)^35^, LMP1 (EBV)^36^, Tax homologs (HTLV-3/4)^37^, vIRF-1/3/4 (KSHV)^38^, E4orf6/7 homologs (HAdv)^39^, and many others. While earlier studies have assayed chromatin-interacting capabilities for a portion of our screened vTR members (chiefly via clonal bulk ChiP-seq and ATAC-seq)^5,40^, genome-wide accessibility changes induced by most variants have not previously been singly characterized.

**Figure 2:**
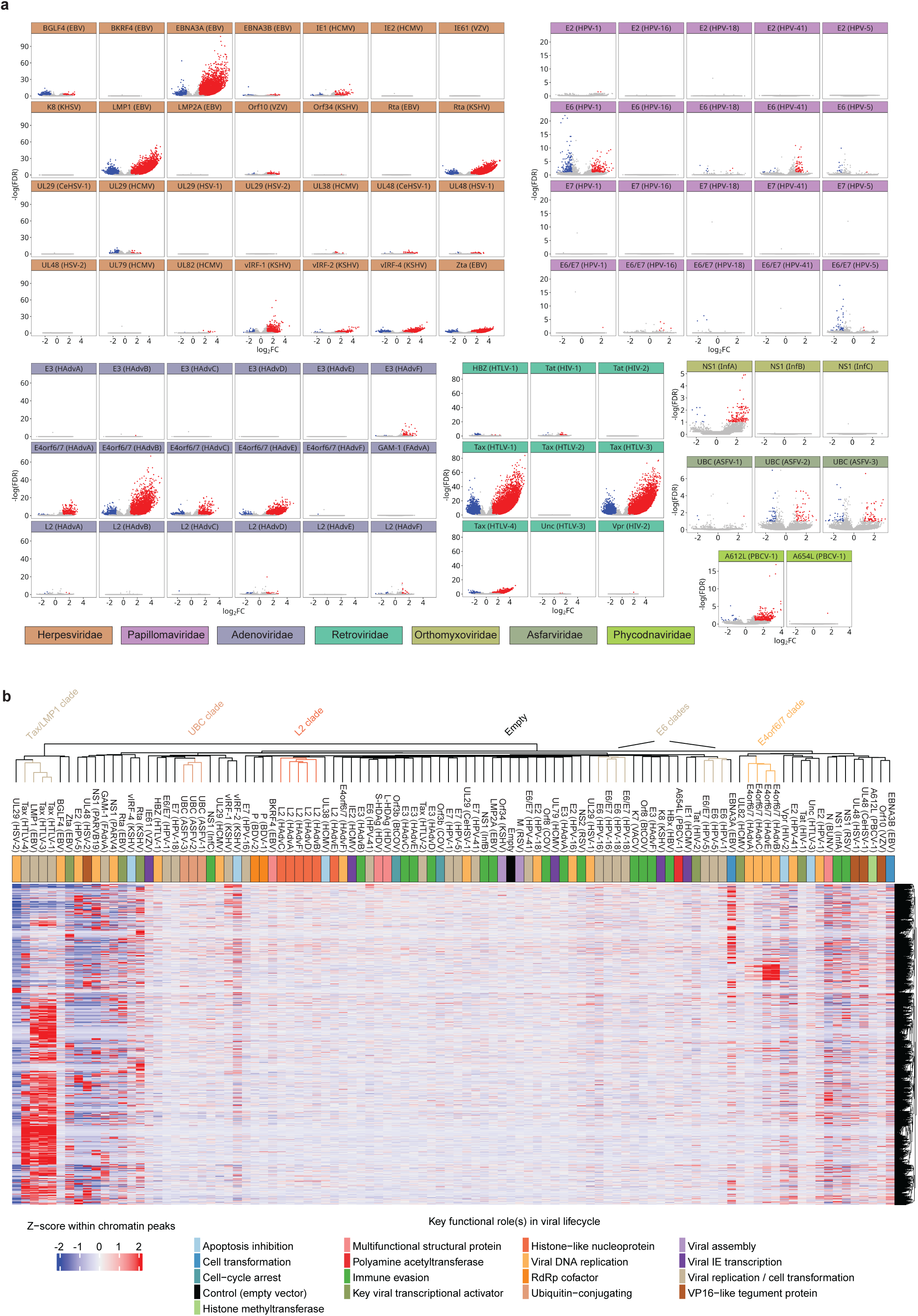
Chromatin dysregulation is a frequently observed feature of vTR-host interactions. **(a)** Volcano plots for vTR variants comparing pseudobulk replicates to EV control, with −log(FDR) versus log_2_FC displayed in each subplot for called peaks. Increased accessibility peaks (FDR ≤ 0.1 and log_2_FC ≥ 1) are colored red, and decreased accessibility peaks (FDR ≤ 0.1 and log_2_FC ≤ −1) are colored blue. Plot headers are colored by viral family. Only families with at least one vTR with > 50 significantly dysregulated peaks are displayed. **(b)** Hierarchically clustered vTRs derived from z-score normalized accessibility at 64,202 peaks that are differentially accessible (FDR ≤ 0.1 and log_2_FC ≥ 1) in at least one vTR. Examples of vTR clades that clustered based on shared chromatin dysregulation patterns are highlighted. Mechanistic metadata for each variant row is displayed which colors each vTR by primary role(s) in the viral infection lifecycle.

The majority of vTR chromatin dysregulation for each variant was measured as increased, rather than decreased, accessibility peaks relative to the EV control. Notable exceptions stood out: BGFL4 (EBV), HBZ (HTLV-1), and UL29 (HCMV) each produced substantially more decreased chromatin accessibility peaks. BGLF4 and HBZ have been identified as potent inhibitors of host NF-κB and AP-1 TF family activities, respectively, which may contribute to the observed preferential compaction over opening of host chromatin sites (a hypothesis which we expand on later)^41,42^. The contributing factors for compaction caused by UL29 are more uncertain, yet one possibility could be the vTR’s previously noted interaction with the host cell Nucleosome Remodeling and Deacetylase (NuRD) complex^43^. The abundance of decreased accessibility peaks noted for many other vTRs, even those expected to primarily act as chromatin pioneering agents, can at least partially be attributed to the redirecting of cell chromatin architectural proteins (ex. SWI/SNF, NuRD, CTCF) or related cofactors due to strong vTR epigenetic modification activities^13^. Abundant opening of large swathes of the typically compacted genome can either displace chromatin-anchored proteins or titrate related architectural proteins from other regions^44^. In a later section, we expand on our hypothesis that much of the decreased accessibility signals reported here are due to non-specific titration or redirection of chromatin structural components, rather than targeted vTR-compacting activities (Supplementary Fig 4).

Many other vTRs produced few to no substantially dysregulated peaks. Multiple factors could contribute to this lack of signal. Firstly, these variants may not exert chromatin dysregulation as a component of their role in their respective viral lifecycles. Additionally, the expression of single vTRs outside their typical physiological context (alongside co-expressed viral proteins in a target host cell) may prevent the detection of certain chromatin interactions. We also included Tax (HTLV-2) as a control, which was expected to produce little to no epigenetic disruption. While three of the assayed Tax homologs (Tax HTLV-1/3/4) have been reported to substantially alter host chromatin, Tax (HTLV-2) lacks a functional nuclear-localization sequence (NLS) and has been shown to produce muted to no chromatin disruption in affected cells^37^. The PROD-ATAC screen recapitulated this expected result, with zero differentially expressed peaks resulting from Tax (HTLV-2) expression compared to the thousands observed for the other Tax homologs. The identification of dozens of vTRs as substantial chromatin remodelers in an infection-agnostic context highlights their singular contributions to host cell epigenetic shifts without reliance upon other viral proteins or cofactors.

We next compared the chromatin dysregulation profiles generated by all 100 library members to each other (as opposed to EV alone) to identify convergent and divergent patterns of vTR-induced epigenetic changes. The chromatin accessibility datasets for all vTRs (and EV) were compared across 60,440 marker peaks (Fig 2b). Variants and peaks were each hierarchically clustered, which illuminated clear relationships based on chromatin disruption patterns across the vTR set. Unambiguous clustering based on shared chromatin accessibility dysregulation emerged for multiple sets of vTR homologs, including the highlighted UBC (ASFV), L2 (HAdv), E6 (HPV), and E4orf6/7 (HAdv) clades (Fig 2b). The clustering of several homolog sets reveals shared chromatin perturbation patterns while highlighting selected homologs which diverge in functionality. Notable examples of variants which diverged from conserved homolog set functionalities included E4orf6/7 (HAdvF) and Tax (HTLV-2) (Fig 2b). We were additionally able to establish functional relationships between structurally and phylogenetically diverse vTR variants. Three functional Tax homologs (HTLV-1/3/4) clustered tightly with LMP1 (EBV). The Tax and LMP1 proteins share very little sequence and structural homology and are sourced from entirely distinct viral families (Retroviridae and Herpesviridae, respectively). Additionally, several distinct vTRs from the same virus or closely related phylogenetic backgrounds clustered together due to similar chromatin-modulating activities. Examples included EBNA3B (EBV) and Orf10 (VZV), as well as vIRF-1 (KSHV) and vIRF-2 (KSHV). Taken together, our large-scale PROD-ATAC screen revealed dozens of vTRs which substantially alter host chromatin and established functional relationships between both phylogenetically similar and diverse variants with convergent patterns of epigenetic disruption.

### Emergent mechanisms of vTRs as host cell epigenetic modulators

Once we identified dozens of vTRs as chromatin accessibility remodelers, we then sought to parse out the mechanistic roles of vTRs inside cells intimated by human TF (hTF) DNA motif enrichment patterns within newly accessible chromatin regions. More than 30 vTRs generated peaks with significant enrichment for at least one hTF motif (Fig 3a). A vTR which preferentially increases accessibility at chromatin regions enriched for certain hTF motifs indicates either direct vTR binding at such sites, or a vTR-driven upregulation of hTF activity. Several enriched motifs supported previously reported vTR – hTF interactions. In KSHV-infected cells, Rta (KSHV) directly interacts with the host RBP-Jκ TF to activate transcription of multiple host and viral genes which promote lytic reactivation^45^. Our screen showed that the increased accessibility peaks for Rta (KSHV) were dramatically enriched for RBP-Jκ binding sites (-log(*P*) = 2247) (Fig 3b). Previous reports also identified the Tax (HTLV-1/3) homologs and LMP1 (EBV) as strong NF-κB TF family activators within infected cells^37,46^. Our results recapitulated these results by demonstrating strong NF-kB motif enrichment for each vTR (Fig 3b). Importantly, these results also provided the first reported evidence that the Tax (HTLV-4) homolog promotes a similar NF-kB-associated epigenetic shift (Fig 3a, c). In addition to their propensities as NF-kB upregulators, peaks opened by the functional Tax homologs and LMP1 were also enriched for Fos binding sites (among other AP-1 TF motifs) (Fig 3c). A recent study surveyed the genome-wide binding and chromatin accessibility profiles of several EBV proteins, including three members of our set: EBNA3A, EBNA3B, and Zta^40^. Our results corroborated their findings, including strong enrichment of the Pbx TF motif in EBNA3A peaks and AP-1 TF motifs (Fos/Jun/Atf) in both the EBNA3B and Zta peak sets (Fig 3b). Overall, these results demonstrate that PROD-ATAC reliably recapitulates expected epigenetic-modifying functionalities for many vTRs in a pooled assay within an infection-agnostic background.

**Figure 3:**
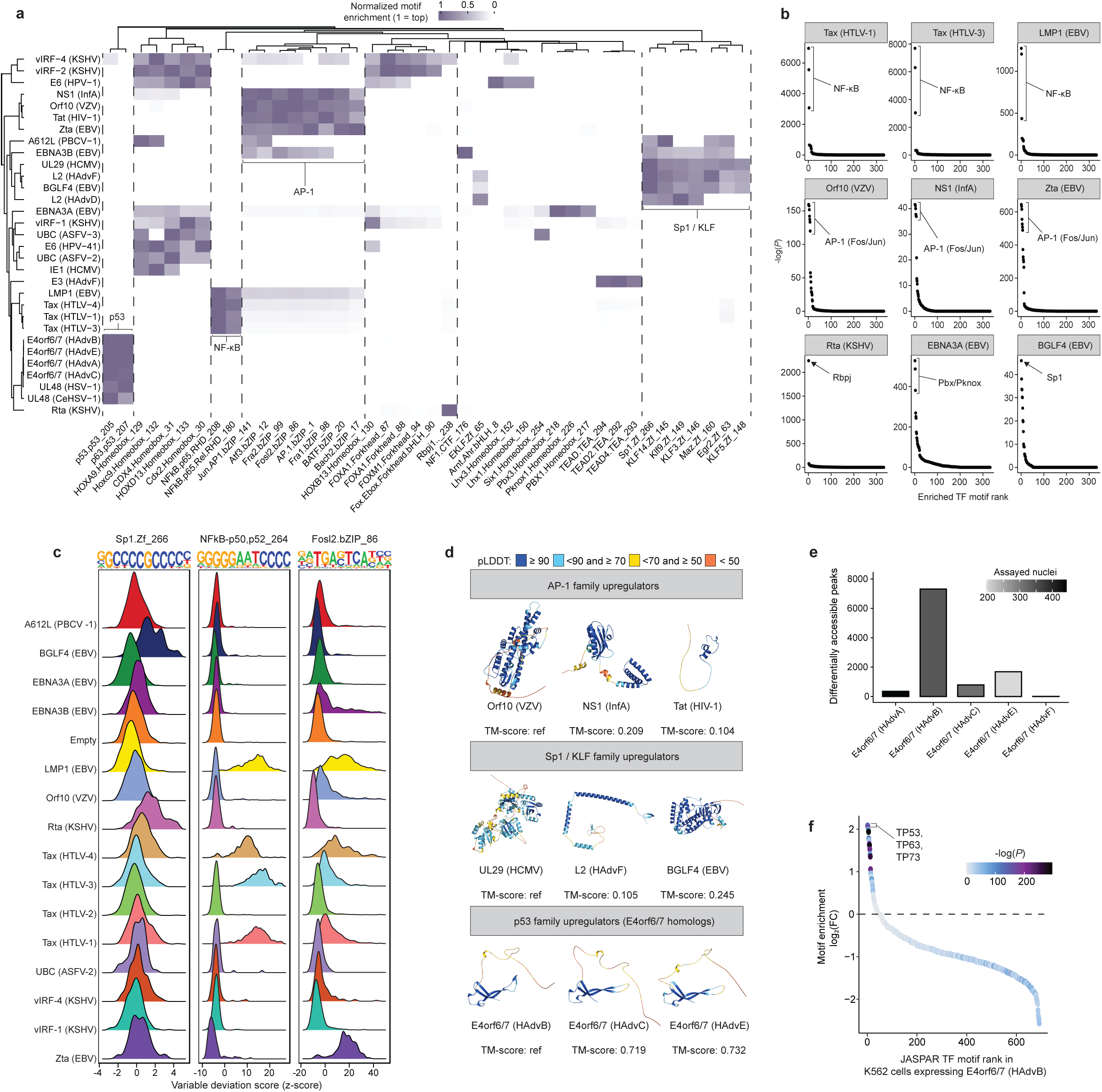
Convergent vTR mechanisms of epigenetic dysregulation. **(a)** Motif enrichment for vTRs with significantly enriched DNA motifs in increased accessibility peaks. Only variants with at least one substantially enriched motif (−log(*P*) > 5)) and only motifs enriched in at least one set of vTR pseudobulk replicates are shown. TF motifs and vTRs are each clustered by Euclidean distance. **(b)** Host cell TF-associated DNA motif enrichment within increased accessibility peaks for nine selected vTRs. Displayed −log(*P*) calculated with hypergeometric tests are plotted versus enrichment ranked order for all queried HOMER motifs. **(c)** Motif deviation plots for a subset of vTRs centered on three distinct TF motifs. Background peaks were identified by sampling peaks based on GC-content similarity and fragment counts across all variants using the Mahalanobis distance (D_M_). Total vTR variant chromVAR deviation (z) scores relative to the background peak set are shown for the selected motifs. **(d)** AlphaFold3 structures of selected vTRs which preferentially enrich distinct host DNA motifs. Positions are colored by predicted Local Distance Difference Test (pLDDT) values. **(e)** Total differentially accessible peaks (FDR ≤ 0.1, log_2_FC ≥ 1 and log_2_FC ≤ −1) for E4orf6/7 homologs. Columns are shaded by number of assayed nuclei. **(f)** JASPAR motif enrichment (log_2_FC) from bulk ATAC-seq of K562 cells expressing E4orf6/7 (HAdvB) relative to K562 cells transduced with empty vector. Points are colored by -log(*P*) motif scores.

Many other vTRs produced motif enrichments which revealed previously unknown direct host DNA or hTF interactions. Several vTRs, including NS1 (InfA) and Orf10 (VZV), were enriched for AP-1 family motifs (Fig 3a). While Tat (HIV-1) and Zta (EBV) are known to increase accessibility at AP-1 binding sites^29,47^, neither NS1 nor Orf10 have been reported as AP-1 activators. Separately, peaks for multiple variants were enriched for Sp1/KLF TF family binding sites. BGLF4 (EBV), UL29 (HCMV), and several L2 (HAdvD/F) homologs were among the Sp1/KLF upregulators (Fig 3a). None of these vTRs were previously known to interact with or modulate the activity of the Sp1/KLF TF family. Interestingly, A612L (PBCV-1) and EBNA3B (EBV) peaks were enriched for both AP-1 and Sp1 motifs (Fig 3a), a finding we emphasize later. Four E4orf6/7 (HAdvA/B/C/E) and two UL48 (HSV-1 and CeHSV-1) variants preferentially opened chromatin enriched for p53 family motifs (Fig 3a). While a single E4orf6/7 homolog has previously been shown to directly interact with host E2F (part of the E2F-Rb-p53 signaling pathway)^39^, we demonstrate specifically that multiple distinct E4orf6/7 homologs serve as p53 family activators based on increased chromatin accessibility in corresponding binding sites genome-wide. The patterns of DNA motif enrichment in peaks opened by vTR activities clearly delineate relationships between distinct variants based on shared mechanisms of epigenetic disruption. We note that many vTRs shown to be activators of AP-1, Sp1/KLF, and p53 TF activity are often highly structurally diverse (Fig 3d). No variants shared substantial predicted folding similarities, apart from the homolog sets (including the E4orf6/7 variants). These results highlight the abundance of viral proteins which, via convergent evolution, assume similar functional roles during infection cycles despite distinct phylogenetic origins. To accurately compare the strengths of chromatin dysregulation and motif enrichment induced by E4orf6/7 variants (and other homolog sets), we can examine the nuclei quantity assigned per variant. We recorded the nuclei captured alongside significant peak counts for the five screened E4orf6/7 homologs. This comparison demonstrated that higher peak count was not solely attributable to larger nuclei populations per homolog, a feature illustrated clearly by the 7,317 peaks (nuclei = 331) for E4orf6/7 (HAdvB) and one peak (nuclei = 324) for E4orf6/7 (HAdvF) (Fig 3e).

The enriched motifs in decreased accessibility peak sets were largely similar across the vTR library. Highly conserved motif enrichment patterns (generally HOX/CDX TF motifs) observed for dozens of vTRs which produced unique enrichments in open chromatin regions support our earlier assertion that a majority of observed compaction is driven by the redirection of chromatin architectural proteins rather than targeted vTR compaction (Supplementary Fig 4). Notable exceptions include the three UBC (ASFV) homologs^48^, HBZ (HTLV-1), UL29 (HCMV), and BGLF4 (EBV), for which the latter three vTRs preferentially closed peaks (as mentioned previously). Closed peaks for the HBZ and the UBC homologs were specifically enriched for AP-1 family motifs. HBZ^42^ and one UBC^49^ variant were previously shown to inhibit AP-1 activity, which these findings corroborate. Interestingly, the enriched AP-1 family motifs for these variants were entirely distinct from the AP-1 motifs observed for the increased accessibility peak sets, which indicates alternative AP-1 complex interactions for the vTR up- and downregulators in our library. BGLF4 did not produce a motif enrichment pattern distinct from most other vTRs, which may denote BGLF4-induced compaction is due to previously reported HDAC1/2 (or other chromatin structural protein) interactions rather than specific hTF dysregulation^50^. The NF1 TF motif was substantially enriched in UL29 decreased accessibility peaks (-log(*P*) = 48), which indicates UL29 downregulates tumor suppressor NF1 activity. This evidence suggests a novel UL29 – NF1 interaction (direct or indirect) inside host cells, which may contribute toward HCMV’s activity as an oncomodulator^51^. Taken together, these results demonstrate that valuable mechanistic inferences can be made into vTR functions inside host cells using both the decreased and increased chromatin accessibility datasets from PROD-ATAC.

Two potential concerns were evident regarding the biological fidelity of the p53 family-activating activities of the E4orf6/7 homologs assayed via PROD-ATAC. The cell type in which we screened these variants, 293T, expresses a portion of the early human adenovirus genome^52^. This region includes the vTRs E1A and E1B, which are critical for the function of other adenoviral proteins but are not confirmed to be necessary for E4orf6/7 activity^53^. Additionally, p53 proteins in 293T cells are sequestered and inhibited by SV40Tag^52^. While E4orf6/7 activation of other highly related p53 family TFs (p63 and p73) could contribute to motif enrichment, SV40Tag activity introduces uncertainty into the biological significance of p53 motif enrichment patterns in the 293T cell context. We addressed both confounding possibilities by performing bulk ATAC-seq on K562 cells which expressed E4orf6/7 (HAdvB). K562 cells do not express any adenovirus proteins and produce a truncated, nonfunctional p53 protein alongside functional p63 and p73^54^. We identified preferential enrichment in p53 family motifs in increased accessibility peaks generated by E4orf6/7 expression in K562 cells relative to the chromatin profile of K562 cells encoding the EV control (Fig 3f). Therefore, E4orf6/7 upregulates the activity of several p53 family members, not exclusively the p53 TF. These findings serve as a controlled set of orthogonal validations which bolster the credibility of the 293T-measured p53 TF family activation induced by several E4orf6/7 variants.

### Comparative epigenetic and transcriptomic analyses reveal shared vTR functional roles

Our analysis of shared chromatin dysregulation and TF DNA motif enrichment patterns established linkages between the broader functional roles of many vTRs inside host cells. We next expanded upon these mechanistic relationships by delving deeply into the individual peak identities and gene expression measurements produced for selected vTRs. We initiated this process by comparing pseudobulk replicate peak profiles for variants we classified as NF-κB and AP-1 TF upregulators. Tax (HTLV-4) produced 3,040 increased accessibility peaks relative to the EV chromatin profile. Yet when we benchmarked Tax (HTLV-4) to Tax (HTLV-1), only 31 significant increased accessibility peaks were measured (a 99% reduction) (Fig 4a). This result indicates that most of the same significantly differentially accessible peaks for Tax (HTLV-4) are similarly dysregulated by Tax (HTLV-1). Comparison to the NF-κB upregulator LMP1 (EBV) also produced a strong (albeit less dramatic) reduction in differential peaks, with >70% of total peaks similarly dysregulated. A separate comparison of Orf10 (VZV) peaks to other AP-1 upregulators likewise established strong peak profile similarities between variants (Fig 4b). The Orf10-EBNA3B comparison yielded a particularly homogenous chromatin landscape, with >80% of all peaks similarly differentially accessible. We can also observe qualitative phenotypic similarities between these variants by visualizing their genotyped nuclei in dimension-reduced space. Highlighted cells which overlap in the shown UMAP representations (derived from Fig 1e) possess structurally similar chromatin dysregulation landscapes. The depicted NF-κB and AP-1 upregulator sets preferentially occupied similar spaces, further demonstrating the profound chromatin profile similarities for these vTRs (Fig 4a, b).

**Figure 4:**
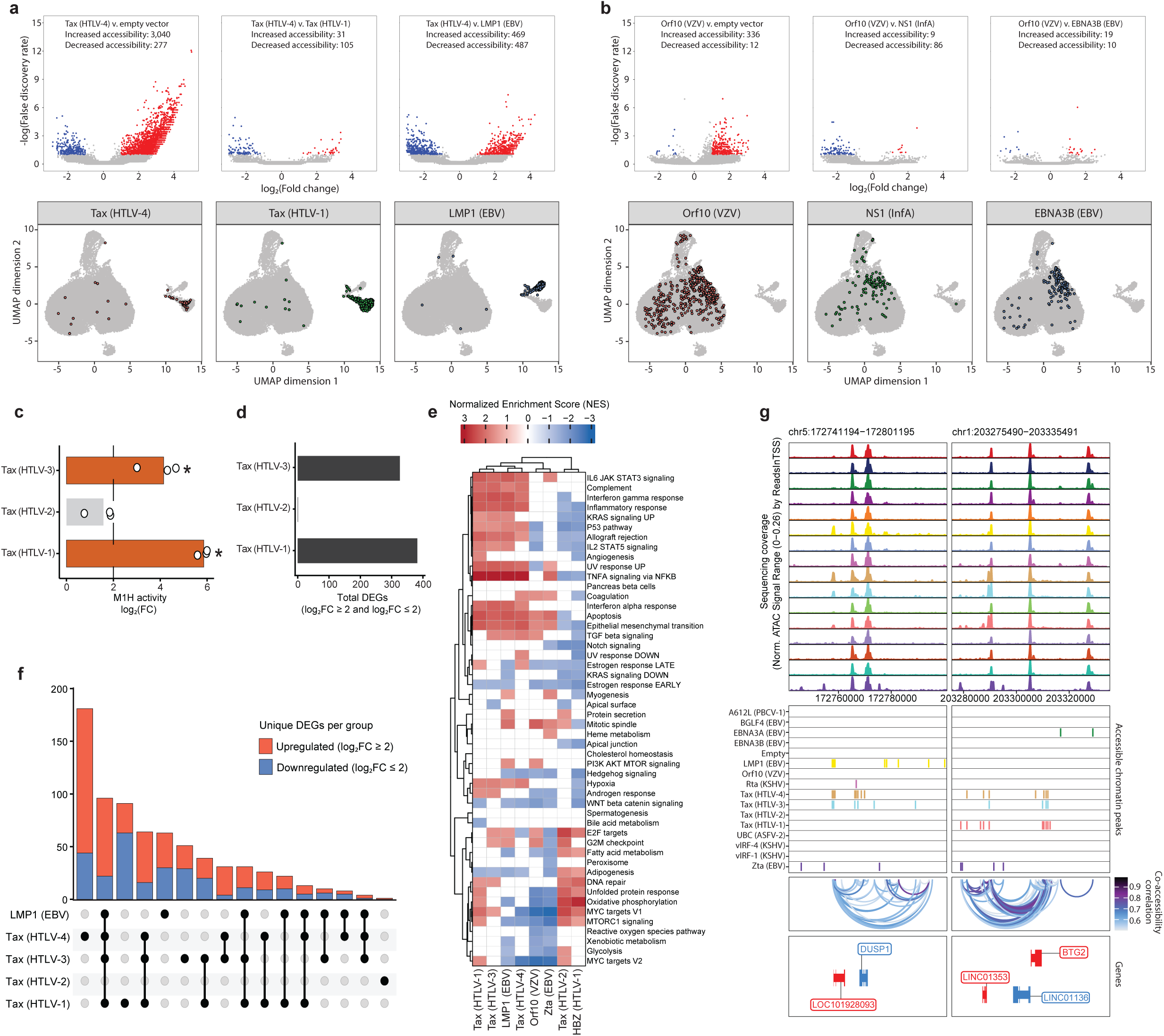
Comparative epigenetic and transcriptomic analyses highlight shared vTR functional roles. **(a)** Volcano plots of comparisons between differentially accessible peaks identified for pseudobulk replicates of Tax (HTLV-4) vs. EV, Tax (HTLV-1), and LMP1 (EBV). As in Fig 2a, plots display −log(FDR) versus log_2_FC. Below are UMAP embeddings post-LSI (as in Fig 1e) with nuclei assigned to the corresponding vTR highlighted. **(b)** Same as (a), now displaying Orf10 (VZV) vs. EV, NS1 (InfA), and EBNA3B (EBV). **(c)** M1H assays with a moderate strength promoter quantified transcriptional activating capabilities of Tax HTLV1-3 homologs in 293T cells, shown as log_2_FC relative to an EV control. **(d)** Bulk RNA-seq results with total host cell differentially expressed genes (DEGs) resulting from Tax HTLV1-3 homolog expression. Significant DEGs were log_2_FC ≥ 2 (upregulated) and log_2_FC ≤ −2 (downregulated). **(e)** Host gene pathway enrichments identified via bulk RNA-seq of vTR-expressing cells. Genes were ranked by log_2_FC and restricted to Hallmark pathways, with non-significant pathways (FDR ≥ 0.05) set to zero and shown as a summary normalized enrichment score (NES) matrix. Pathways which vTR variants upregulated (NES ≥ 1) are shown as red, while downregulated pathways (NES ≤ −1) are shown as blue. **(f)** Unique DEGs associated with each shown combination of LMP1 and Tax HTLV1-4 homologs. **(g)** Genome browser tracks displaying sequencing coverage, significant chromatin peaks (FDR ≤ 0.1 and log_2_FC ≥ 1), and co-accessibility correlation values for vTR peaks in the 60-kb windows shown.

Comparative transcriptomic analyses were performed for a subset of the vTR library to elucidate variant influences on cell multiomic profiles and further highlight mechanistic relationships between vTRs. Previous studies have shown many vTR domains and full-length proteins exhibit strong transcriptional-activating capabilities^7,55^. A moderate-strength promoter mammalian one-hybrid (M1H) system was applied to quantify the transactivating capabilities of several Tax homologs in 293T cells. Tax (HTLV-1) (log_2_FC = 5.86) and Tax (HTLV-3) (log_2_FC = 4.15) were revealed to be strong transactivators, while Tax (HTLV-2) (log_2_FC = 1.58) demonstrated only muted transactivating capability relative to the control (Fig 4c). Next, we generated gene expression datasets via bulk 3’ RNA-seq for 293T-LP cells clonally expressing nine library members (including EV). Each vTR produced significant gene expression dysregulation, with hundreds of differentially expressed genes (DEGs) measured for the three functional Tax (HTLV-1/3/4) homologs, LMP1, and Zta. Only one gene was significantly dysregulated by Tax (HTLV-2), a finding consistent with the vTR’s negligible signals in our ATAC-seq and M1H datasets (Fig 4d). As many vTRs are either themselves oncogenic proteins or are derived from oncoviruses, we performed pathway enrichment analysis on the vTR-induced gene expression shifts centered on the cancer-associated Hallmark gene sets^56^ (Fig 4e). Hierarchical clustering of the vTRs established three distinct groups: NF-κB activators (Tax (HTLV-1/3/4) and LMP1); AP-1 activators (Orf10 and Zta); Tax (HTLV-2) and HBZ. Significant enrichment of multiple pathways was observed for each variant. We particularly note the strong enrichment of interferon and other immune response-associated gene sets by the NF-κB upregulators, and the downregulation of several metabolic and cell growth gene sets (including Myc targets V1/V2) by the AP-1 upregulators.

We further examined the DEG profiles of the NF-κB-activating vTRs (LMP1 and the Tax homologs) to identify unique gene expression changes that might define subtle mechanistic differences between the roles of these variants within cells. Comparative analysis of the DEGs unique to each possible grouping of the NF-κB upregulators revealed that Tax (HTLV-4) alone produced by far the most unique DEGs, with more than 150 DEGs not observed for the other vTRs (Fig 4f). This indicates there are substantial functional differences and distinct cellular interactions for the understudied Tax (HTLV-4) homolog relative to the comparatively well-characterized Tax (HTLV-1/3) homologs^37^, despite each variant sharing largely similar chromatin dysregulation landscapes. LMP1 yielded nearly 100 DEGs also shared with the functional Tax (HTLV-1/3/4) homologs (Fig 4f), which corroborated our previously noted functional similarities between the vTRs. We next demonstrated the utility of our multiomic (epigenetic and transcriptomic) vTR datasets by examining PROD-ATAC genome browser tracks centered at two genes that were uniquely dysregulated by selected variants: DUSP1 (MAPK signaling role)^57^ and BTG2 (tumor suppressor role)^58^ (Fig 4g). RNA-seq revealed that DUSP1 expression was highly enriched by the Tax (HTLV-3/4) homologs (log_2_FC > 3.5 for each), while LMP1 and Tax (HTLV-1) produced no significant DUSP1 expression shift. The Tax (HTLV-3/4) homologs generate accessible peaks immediately up- and downstream of DUSP1, while LMP1 and Tax (HTLV-1) produced no peaks <10 kb upstream of the gene-coding region (Fig 4g). This suggests epigenetic perturbations such as increased promoter and enhancer access are associated with the DUSP1 expression change. Separately, all NF-κB activators strongly increased BTG2 expression (log_2_FC > 4) except for LMP1. The vTRs that enriched BTG2 expression (Tax (HTLV-1/3/4)) opened the chromatin immediately within and surrounding the gene, while LMP1 generated no significant chromatin accessibility within >30 kb of the gene-coding region (Fig 4g). This finding provides yet another direct linkage between vTR-induced chromatin perturbations and dramatic gene expression shifts. Overall, we demonstrate that the complementary inputs of epigenetic and transcriptomic datasets bolster the credibility of both signal types^59^ and illuminate subtle differences that define unique biological roles for otherwise functionally related variants, such as LMP1 and the Tax homologs.

### PROD-ATAC drug-based perturbation screening yields mechanistic insights into avenues of vTR epigenetic modification

No study to date has performed drug-based perturbation screening on pooled protein-coding variants to yield ATAC-seq readouts. We demonstrated the adaptability of our platform by performing PROD-ATAC on a condensed 15-member vTR library in the presence of two separate small molecule inhibitors: the MEK1/2 inhibitor TAK-733^60^, and the BRD inhibitor JQ1^61^. We also screened the same library with a vehicle control (0.1% DMSO) to enable accurate comparisons between treatments. Prior to peak calling and motif enrichment analysis, the nuclei in these three conditions (vehicle control (V.C.), TAK-733, and JQ1) were randomly downsampled to the lowest value per vTR in any condition^62^. For example, 350, 332, and 320 BGLF4 nuclei were genotyped in the TAK-733, JQ1, and control treatments, respectively. We therefore randomly downsampled the TAK-733 and JQ1 samples to match the 320 BGLF4 nuclei assayed in the control condition. Each drug treatment was selected to target distinct cell regulatory features. MEK1/2 inhibition by TAK-733 disrupts MAPK signaling cascades^60^, while JQ1 inhibits the chromatin interaction of BRD4, a key scaffold and coactivator protein^61^. These small molecule perturbations provided separate insights into the functional roles of these vTRs inside host cells.

We first measured the drug-induced shifts (log_2_FC) in peaks opened by vTRs relative to the control condition (Fig 5a). Several vTR-chromatin pioneering activities were dramatically reduced by TAK-733. Opened peak profiles for Orf10 and EBNA3B, which we previously noted to produce highly similar chromatin landscapes, were the most impacted with log_2_FC values of - 3.84 and -2.26 for Orf10 and EBNA3B, respectively (Fig 5a). Chromatin opening activities for three other vTRs were also significantly downregulated: A612L (log_2_FC = -1.72), vIRF-4 (log_2_FC = -1.42), and BGFL4 (log_2_FC = -1.95) (Fig 5a). Each vTR inhibited by TAK-733 increased accessibility in chromatin regions enriched for AP-1 motifs, with the sole exception of BGLF4. We note that BGLF4, A612L, and EBNA3B were all shown by the large-scale screen and our condensed library control to produce significant AP-1 and Sp1/KLF TF motif enrichment. As these five AP-1 / Sp1 – upregulating vTRs were specifically inhibited by TAK-733, this data indicates that MEK1/2 activity (and the associated MAPK signaling pathway) is directly involved in the epigenetic dysregulation induced by these variants. Separately, the JQ1 treatment substantially decreased chromatin-opening by EBNA3B (log_2_FC = -3.01) and vIRF-1 (log_2_FC = - 2.51) (Fig 5a). While TAK-733 treatment solely decreased accessible peak counts, JQ1 also increased (log_2_FC > 0.5) the chromatin-opening activities for three vTRs: A612L, Orf10, and Tax (HTLV-4).

**Figure 5:**
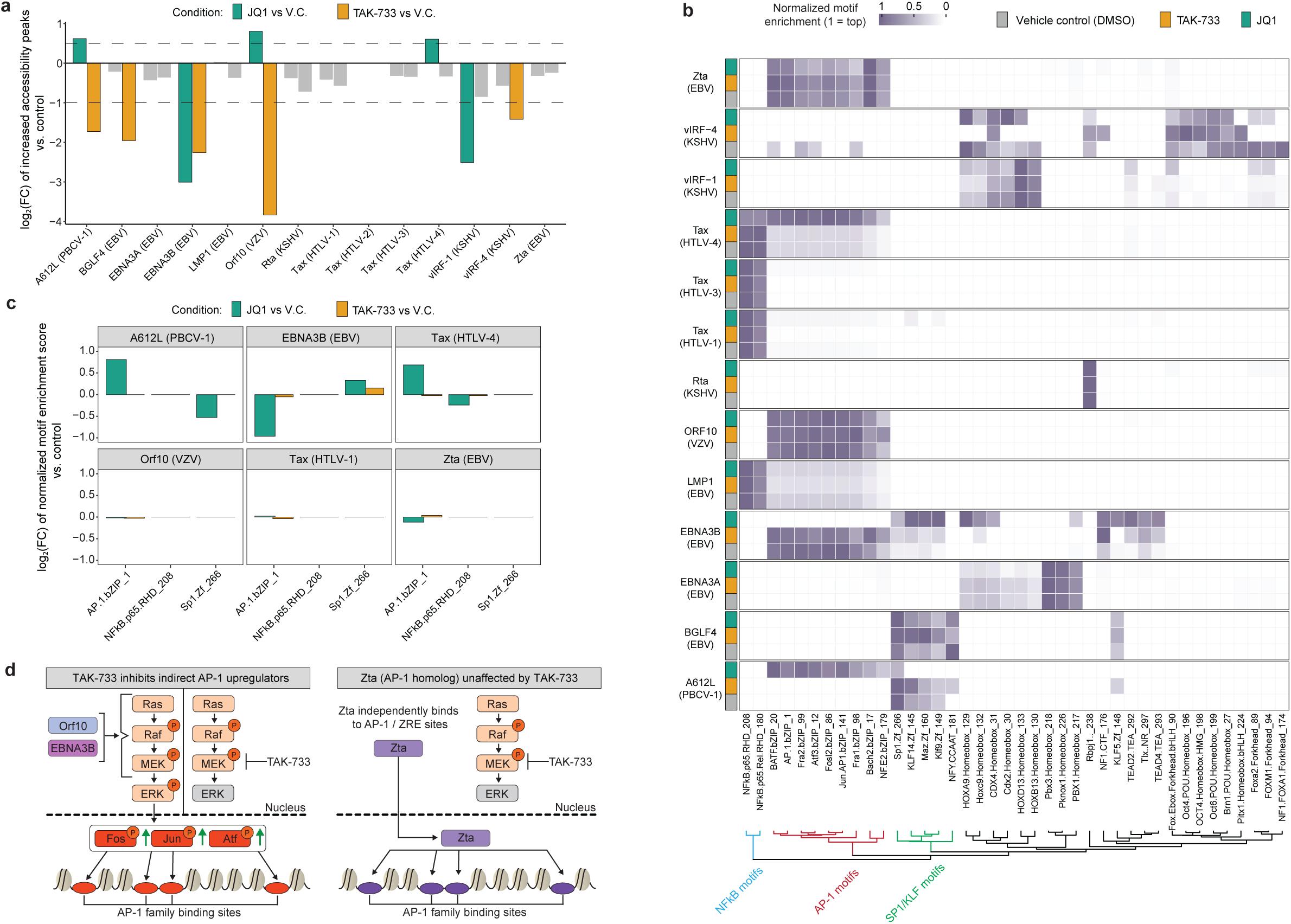
PROD-ATAC drug-based perturbations reveal vTR host-dependencies. **(a)** Shifts in vTR increased accessibility peak counts (log_2_FC) in drug-treated conditions relative to the vehicle control. **(b)** DNA motif enrichment patterns for vTRs in control and treated conditions. Motifs are clustered by Euclidean distance. Only motifs which were substantially enriched in at least one set of vTR pseudobulk replicates (−log(*P*) > 2)) are displayed. **(c)** Drug-induced shifts in representative motifs (log_2_FC) of values from Fig 5b) relative to the control. **(d)** Schematic depicting specific inhibitory activity of TAK-733 against vTRs reliant on MAPK/ERK pathway for inducing host chromatin dysregulation.

We next examined shifts in vTR motif enrichment patterns which resulted from JQ1 activity (Fig 5b). JQ1 treatment altered the DNA motif enrichment pattern in accessible regions for three vTRs: A612L, Tax (HTLV-4), and EBNA3B. A612L and Tax (HTLV-4) accessible peaks were primarily enriched for Sp1/KLF and NF-κB motifs in the control condition. Upon JQ1 treatment, open chromatin regions for both vTRs were predominantly enriched for AP-1 family motifs (Fig 5c). In contrast, JQ1 activity shifted the AP-1 motif dominated profile of EBNA3B peaks to an overrepresentation of Sp1/KLF TF motifs. Interestingly, the vTRs with increased AP-1 motif representation, A612L and Tax (HTLV-4), generated >40% more open peaks with JQ1 treatment (Fig 5a). EBNA3B treated with JQ1 produced ∼88% fewer peaks and the remaining open sites were enriched for Sp1/KLF motifs. We next compared the normalized motif enrichment scores of representative motifs in the AP-1, Sp1, and NF-κB families for these three vTRs when treated with JQ1 (Fig 5c). Proportional abundance of a representative AP-1 motif in open peaks was >50% increased for A612L and Tax (HTLV-4) and >50% decreased for EBNA3B. Several other vTRs produced no substantial motif abundance shifts in the JQ1 condition. This includes Orf10, which also produced a >40% peak count increase with JQ1 treatment. Additionally, vIRF-1 opened significantly fewer peaks in the JQ1 condition without a concomitant TF motif shift.

Taken together, the TF motif enrichments and chromatin-opening strengths altered by JQ1 provide several mechanistic insights into the epigenome-modifying features of the assayed vTRs. First, three vTRs (A612L, Tax (HTLV-4), and Orf10) all opened a significant number of additional chromatin regions. Two of these vTRs, A612L and Tax (HTLV-4), shared a preferential shift toward AP-1 TF motif enrichment. BRD4 inhibition by JQ1 has been shown to alter the topology of the host genome, preferentially closing and opening access to various super-enhancer sites and key regulatory features^63^. The shared increase in chromatin-pioneering strength and DNA motif shifts for these variants indicate that each vTR interacts with the same or functionally related host cell factors that pioneer regions made newly accessible by JQ1 remodeling of the genomic architecture. As the majority of Orf10 peaks in the control were already enriched for AP-1 motifs, generating more peaks with similar AP-1 sites in the JQ1 treatment would not increase AP-1 motif proportional abundance. Second, the EBNA3B AP-1 motif peaks were significantly downregulated by JQ1 treatment. Therefore, EBNA3B-induced AP-1 upregulation is at least partially dependent on host factors or cell signaling pathways which are not strictly required for pioneering at AP-1 sites by A612L, Tax (HTLV-4), and Orf10.

TAK-733 treatment revealed cell signaling pathways which are critical for vTR-induced chromatin dysregulation. The full-scale PROD-ATAC screen identified several vTRs which upregulate AP-1 activity, including EBNA3B, Orf10, and Zta. TAK-733 treatment dramatically reduced the pioneering at AP-1 sites facilitated by EBNA3B and Orf10, yet produced no significant change in Zta-induced chromatin dysregulation (Fig 5a). Zta acts as a functional AP-1 homolog and has previously been shown to pioneer AP-1 and Zta recognition element (ZRE) sites in host cell chromatin^29^. EBNA3B and Orf10, however, have not been shown to independently bind host DNA and instead indirectly induce chromatin dysregulation through interactions with host TFs and other regulatory features^40,64^. TAK-733 allosterically inhibits MEK1/2 activity, thereby interrupting the MAPK (Ras-Raf-MEK-ERK) phosphorylation cascade and preventing AP-1 TF upregulation^60^ (Fig 5d). As EBNA3B and Orf10 were specifically inhibited by TAK-733, the chromatin-dysregulation mechanisms of these vTRs are highly dependent on MAPK signaling. In contrast, Zta-induced chromatin pioneering is not inhibited by TAK-733 as the vTR binds directly to AP-1 / ZRE sites. Therefore, AP-1 motif enrichment in peaks opened by Zta is largely MEK1/2 independent (Fig 5d). Overall, the PROD-ATAC drug perturbation screens provided valuable insights into the mechanisms of chromatin pioneering by vTRs and identified cell signaling pathways and regulatory features that were previously unknown to be critical for vTR-induced epigenetic dysregulation inside host cells.

### PROD-ATAC analyses of vTR-induced chromatin disruptions are robust and reproducible

In a previous study, we demonstrated that PROD-ATAC single-cell datasets are highly reproducible and closely reflect key features of chromatin accessibility landscapes generated by clonal bulk ATAC assays^23^. That initial work, which centered on oncofusion-induced chromatin dysregulation, revealed PROD-ATAC peak accessibility profiles strongly correlate with bulk ATAC landscapes that demand at least 100,000 cells (for duplicate assays). We applied the same quality control metrics here to ensure that our full-scale and drug perturbation vTR PROD-ATAC screens also yielded similarly robust epigenetic dysregulation datasets. First, selected variants were recombined into 293T-LP (identical to the library assembly workflow) to produce clonal vTR-expressing cell lines. These clones were then induced in the same manner as the vTR library assays and duplicate samples for each variant were harvested for bulk ATAC analysis. Second, we compared the chromatin landscapes for selected vTRs screened in the large-scale assay (replicate 1) and condensed vTR library control condition (replicate 2). As in the oncofusion study, we then quantified correlation between the variants in each assay format at dynamic sites defined by vTR chromatin profiles from the large-scale experiment.

Comparison of the peak accessibility scores at dynamic sites for the two PROD-ATAC replicates and bulk ATAC samples revealed remarkably high correlation by genotype for each assay type (Fig 6a). As expected, hierarchical clustering of the variant correlation values also clearly delineated the distinct epigenetic dysregulation phenotypes observed by LMP1 and Tax (HTLV-1) (NF-κB upregulators) and their substantial dissimilarity to Zta (AP-1 upregulator). Overall, this analysis demonstrated that PROD-ATAC datasets are reproducible and recapitulate the chromatin landscapes generated by arrayed bulk ATAC sequencing efforts which use >10,000x the number of nuclei assayed per variant in our pooled scATAC platform.

**Figure 6:**
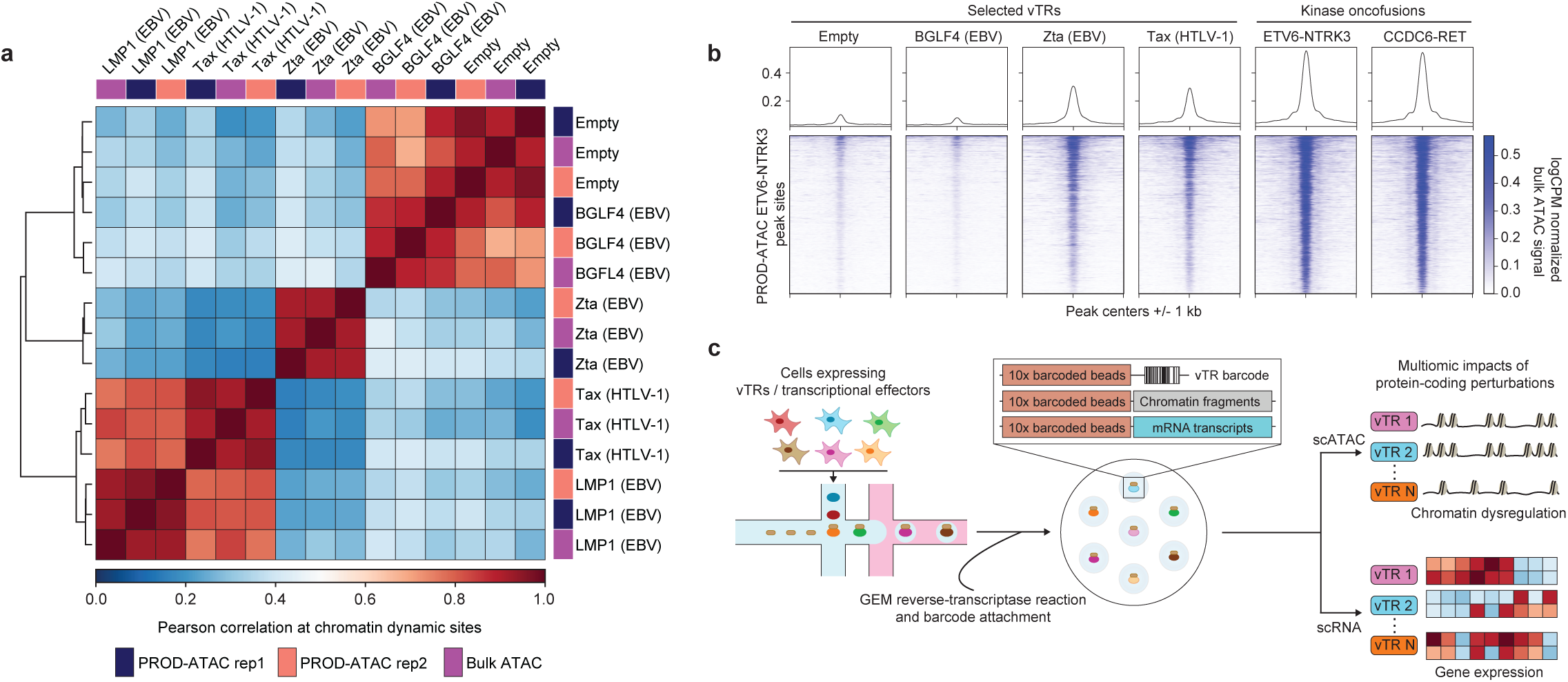
PROD-ATAC chromatin dysregulation measurements are robust and illuminate functional similarities across distinct protein libraries. **(a)** Hierarchically clustered Pearson correlation at dynamic sites for bulk ATAC chromatin profiles compared to those generated by PROD-ATAC replicate experiments. Dynamic sites were defined as differentially accessible peaks in replicate 1 (large-scale PROD-ATAC assay). **(b)** Metaplots and tornado plots for differentially accessible sites identified from pseudobulk replicates of ETV6-NTRK3 generated by PROD-ATAC. Plots depict log(counts per million) (logCPM) normalized ATAC signals from bulk ATAC replicates of cells expressing the shown vTRs and oncofusions. **(c)** Future adaptations of the PROD-ATAC platform will emphasize generating multiomic profiles from the same cells to facilitate mechanistic inferences only feasible via transcriptome-epigenome linkages.

In our initial PROD-ATAC study, we also discovered that many oncofusion (cancer-causing fusion) proteins, including ETV6-NTRK3^65^ and CCDC6-RET^66^, opened thousands of peaks which were enriched for AP-1 motifs^23^. These oncofusions, which drive malignancies including lung adenocarcinoma and non-small cell lung cancer, each contain a functional kinase domain which facilitates the upregulation of host MAPK signaling and AP-1 TF activity. We note that peaks produced by several vTRs in our screen, including Zta and Tax (HTLV-1), partially overlap with the chromatin regions dysregulated by these kinase fusions (Fig 6b). As we highlighted in an earlier section, both Zta and several Tax homologs opened chromatin regions enriched for AP-1 TF binding sites. Notably, Zta and Tax are derived from oncoviruses (EBV and HTLV, respectively) and are themselves characterized as oncogenic proteins^29,37^. Expanded studies to examine similar chromatin dysregulation patterns across vTRs, oncofusions, and related epigenome dysregulators may reveal mechanisms by which oncoviruses and their associated vTRs function as oncogenic modulators.

## Discussion

The single-cell PROD-ATAC assays performed in this study constitute the first systematic surveys of host chromatin dysregulation induced by viral proteins in pooled formats. Our full-scale screen generated chromatin accessibility landscapes for more than 100 vTRs sourced from fifteen viral families. The resulting dataset recapitulated chromatin dysregulation known to be driven by well-characterized vTRs, including Zta and EBNA3A from EBV. Simultaneously, many more vTRs without a reported capability to alter host cell chromatin accessibility were identified as potent chromatin dysregulators, including Orf10 (VZV) and BGLF4 (EBV). The scale of this experiment highlights the advantages of pooled epigenomic screening for functionally characterizing viral proteins. To reproduce the scale of epigenetic disruptions captured by the initial PROD-ATAC assay, an arrayed bulk ATAC screening effort would have required over 100 cell lines and more than 200 NGS library preparations. The full-scale PROD-ATAC assay required a single flask of pooled vTR-expressing 293T-LP cells and one 10x Genomics EpiATAC chip to generate chromatin landscapes for over 100 vTRs. Furthermore, the pooled format of the PROD-ATAC assay provides unique benefits over arrayed ATAC experiments. The shared cell background and singular genome transposition event negate the sample-to-sample variations and batch effects which may result from arrayed bulk ATAC assays that obfuscate meaningful comparisons between vTR chromatin dysregulation phenotypes^67^.

Our analyses reveal that host chromatin remodeling is a widespread and often heterogeneous property of vTRs. Approximately one-third of the screened vTRs significantly dysregulated host chromatin. Effects ranged from subtle local perturbations to dramatic genome-wide remodeling at thousands to tens of thousands of loci. These activities were distributed across multiple viral families and genome architectures, suggesting that epigenetic manipulation of host chromatin represents a broadly conserved strategy employed during viral infection lifecycles^5,8^. While the full-scale PROD-ATAC assay screened over 100 variants, this represents a fraction of the many thousands of identified and hypothetical vTRs (including orthologs)^5,68^. The identification of several previously unrecognized chromatin modulators from our sampling of the massive vTR sequence space suggests that viral proteomes boast a far richer repertoire of host chromatin-interacting functions than can be deciphered by targeted studies of individual viral proteins.

Comparative analyses of the accessibility landscapes generated by diverse vTRs revealed striking similarities in mechanisms of chromatin dysregulation. Chromatin accessibility landscapes were shared not solely among vTR homologs but also across phylogenetically and structurally unrelated variants. For example, similar accessibility profiles were generated by EBNA3B (EBV) and Orf10 (VZV) despite their low sequence similarity and distinct viral origins. Mechanistic classification of vTRs based on TF motif enrichment in accessible chromatin regions highlighted that diverse vTRs often interact with similar host regulatory features. Regions opened by several variants were separately enriched for p53, AP-1, NF-κB, and Sp1/KLF TF family binding motifs, all of which play central roles in cellular stress responses, immune signaling, and proliferative control. Structurally unrelated vTRs from diverse viral lineages frequently targeted the same TF networks. For instance, Tat (HIV-1), NS1 (InfA), and Orf10 (VZV) all preferentially opened chromatin regions enriched for AP-1 motifs despite their disparate structures and evolutionary origins. These findings suggest that diverse viruses often evolve similar functionalities to engage key host signaling pathways and transcriptional hubs which broadly reprogram cell states.

The functional relationships of several vTRs were further elucidated by comparing individual chromatin peak similarities and host transcriptome profiles. Variants which upregulated the same TF families frequently perturbed identical chromatin loci. For instance, pairwise comparison of the peaks opened by the NF-κB upregulators Tax (HTLV-4) and LMP1 (EBV) revealed that most sites dysregulated by Tax (HTLV-4) are similarly dysregulated by LMP1. Separately, our integration of chromatin accessibility and transcriptomic datasets demonstrated that epigenetic similarities often translate into overlapping transcriptome perturbations. Bulk RNA-seq analyses revealed that several Tax homologs and LMP1 induced hundreds of shared differentially expressed genes, consistent with their similar chromatin landscapes. Conversely, subtle differences in chromatin dysregulation by related vTRs were also associated with distinct transcriptional profiles. Notably, the understudied Tax (HTLV-4) homolog generated over 150 differentially expressed genes not dysregulated by other Tax proteins or LMP1. These multiomic analyses provided rich insights into the functional relationships of vTRs as host epigenome dysregulators.

We improved the PROD-ATAC platform by implementing a drug-perturbation workflow to screen a condensed vTR library treated with small molecule inhibitors. Screening vTRs in the presence of the MEK inhibitor TAK-733 and the BET inhibitor JQ1 generated a set of drug-dependent chromatin accessibility landscapes that revealed key host regulatory dependencies. MAPK inhibition by TAK-733 strongly suppressed chromatin remodeling induced by several AP-1-associated vTRs, indicating that their epigenetic activity relies on host signaling cascades that activate AP-1 TFs. In contrast, Zta-induced chromatin dysregulation was insensitive to MAPK inhibition, consistent with its ability to independently bind AP-1 / ZRE elements. BET inhibition by JQ1 shifted TF motif enrichment patterns for several vTRs, suggesting that chromatin scaffolding proteins such as BRD4 shape the genomic regions targeted by vTRs. These drug perturbation experiments demonstrate that PROD-ATAC pooled epigenomic screening can be adapted to identify host features required for viral epigenetic dysregulation at scales that are impractical using traditional arrayed approaches. More broadly, the ability to perform pooled drug perturbation screens with chromatin accessibility readouts opens new avenues for systematically identifying host dependencies of vTRs and other protein-coding variant libraries, such as cancer fusion proteins and artificial transcription factors.

In summary, this study provides the most comprehensive survey to date of host epigenetic dysregulation induced by viral proteins. Our combined full-scale and drug-perturbation PROD-ATAC assays together produced over 150 unique chromatin accessibility landscapes shaped by vTRs. These datasets reveal convergent mechanisms through which vTRs reprogram host regulatory networks and establish a scalable framework for systematically dissecting virus-host chromatin interactions. Ongoing development of multiomic implementations of our pooled single-cell platform (Fig 6c), integrating same-cell transcriptomic and chromatin accessibility profiles, will enable the characterization of coordinated transcriptional and epigenetic programs driven by vTRs and other protein-coding libraries.

## Acknowledgements

J.E.C. was supported through the William H. Peterson Fellowship, awarded by the University of Wisconsin-Department of Biochemistry. This work was funded in part through the S.C. Fang Professorship and Vilas Associate awards, both granted to S.R.. Additional funding was provided through the NIH grant R35 GM128625, awarded to J.F.B.. We thank the University of Wisconsin-Madison Center for Genomic Science Innovation for providing the 10x Genomics Chromium X. Most DNA amplicon sequencing runs and TapeStation analyses were performed at the University of Wisconsin-Madison Biotechnology Center’s DNA Sequencing Facility (RRID: SCR_017759). Microscopy was performed at the University of Wisconsin-Madison Biochemistry Optical Core, which was established through support from the University of Wisconsin-Madison Department of Biochemistry Endowment.

## Ethics declaration

The authors declare no competing interests.

## Methods

### Cell culture

The 293T-LP clonal line was used for creating and screening all PROD-ATAC libraries in this study. We previously described in depth the engineering of the 293T-LP cell line to function as a landing pad for PROD-ATAC protein-coding variant libraries^23^. 293T and 293T-LP cells were cultured at 37 °C and 5% CO2 in high-glucose DMEM supplemented with 10% Tet System Approved FBS (Gibco), 1% penicillin–streptomycin and 1% GlutaMAX (Gibco). K562 cells were purchased from the American Type Culture Collection. K562 cells were cultured at 37 °C and 5% CO_2_ in RPMI supplemented with 10% Tet System Approved FBS (Gibco), 1% penicillin–streptomycin and 1% GlutaMAX (Gibco).

Cells were regularly confirmed to be mycoplasma free by testing with the Venor GeM Mycoplasma Detection Kit (Millipore Sigma). All 293T and K562 cell stocks were stored in complete media with 5% v/v sterile DMSO in liquid nitrogen vapor phase.

### PROD-ATAC variant barcoding design

The vTR variants were barcoded exactly as described in our previous oncofusion-centered PROD-ATAC publication^23^. Each vTR library member was designed to encode two barcodes. The first barcode is a constant 12-nucleotide (nt) DNA sequence synthesized alongside each vTR coding sequence. The DNABarcodes R package was used to create 200 12-nt GC balanced barcodes with a minimum Hamming distance of 5. Each member of the variant library was randomly assigned to one 12-nt barcode. Ideally, at this stage Nextera adapter sequences would be included in the synthesized vTR sequences, which would make the single 12-nt barcode sufficient for variant genotyping. However, commercial oligonucleotide synthesis companies cannot typically generate sequences encoding Nextera or TruSeq adapters internally as they are also used for NGS quality control prior to shipment. To enable variant genotyping during scATAC library generation, we inserted a second barcode of 16 random nucleotides (N16) between Nextera adapters to a position adjacent to the 12-nt constant barcode. This configuration situates the 12-nt and random barcodes within 113 bp of each other. This enabled sequencing of the constant 12-nt and N16 barcodes together via short read sequencing to map the twin variant barcodes to their corresponding vTR genotype. We also added the primer binding sequences necessary for PROD-ATAC and Spear-ATAC assays: the oSP1735 site (GCTTACATTTTACATGATAGGCTTGG), which allows for in-droplet exponential barcode amplification; the oSP2053 site (AAGTATCCCTTGGAGAACCACCTTG), which enables linear amplification with a biotinylated primer. These sequences together enable the generation of a barcode library alongside the scATAC-seq libraries which is used to genotype each nucleus.

### Donor plasmid library construction

All 110 library member sequences (109 vTRs + EV control) were cloned via BsmBI Golden Gate Assembly into an attB-containing donor plasmid to later be recombined into attP-containing 293T-LP cells. Each vTR sequence was codon-normalized for human expression using Twist Bioscience online software. Internal BsmBI sites were removed from each sequence by creating synonymous substitutions that avoided rare codons. We added the 12-nt barcodes (described above) and universal primer binding sites such that all variants could be amplified with a single primer pair. This primer pair included part of the Nextera adapter sequences and the oSP2053 primer binding site. The primers also encoded the BsmBI sites needed for downstream Golden Gate Assembly. All variant sequences were ordered from Twist Bioscience. PCR amplification of each library member yielded one clearly dominant product per variant at the correct size by agarose electrophoresis. PCR products for these variants were quantified using an AccuClear Ultra High Sensitivity dsDNA Quantitation Kit and pooled to target equimolar distributions. The one exception to this pooling ratio was that we doubled the amount of EV DNA to slightly bias EV nuclei capture during scATAC-seq. This ensured that we could establish an empirical null distribution to which vTR signals could be compared.

The attB-containing backbone was linearized by PCR to be joined with the donor sequences (above) for BsmBI Golden Gate Assembly. This PCR added the N16 barcode, part of the Nextera adapters, and BsmBI cut sites. Following amplification, the backbone PCR reactions were DpnI digested by spiking in 2 µL of DpnI (20 U/µL), 10 µL of NEB 10x rCutSmart buffer and water to reach a total volume of 100 µL. The reaction proceeded at 37 °C for 1 hr, followed by a 20-min inactivation at 80 °C before being cleaned up. All PCR-amplified samples were cleaned up prior to assembly steps (Omega Bio-Tek, CP buffer / DNA wash buffer). A single Golden Gate Assembly reaction was then performed to insert the pool of variant members into the attB-containing backbone. This reaction combined 150 ng of backbone with the pooled vTR library PCR product for an approximately 2:1 backbone:insert molar ratio. To this was added 2 µl of 10x T4 ligase buffer, 1 µl of BsmBI-v2 and water to bring the total reaction to 20 µL. The reaction proceeded at 42 °C for 1 hr, followed by 60 °C for 5 min. The 20 µl product was dialyzed on an MF-Millipore Membrane Filter (0.025-µm pore size) for 1 hr in nuclease-free water.

Finally, we created a bacterial stock of the library. First, 2 µl of the dialyzed Golden Gate product was transformed into 100 µl of electrocompetent DH10B *E. coli* cells using a Bio-Rad MicroPulser (165-2100) on the Ec2 setting (2-mm cuvette, 2.5 kV). Cells were immediately resuspended in 973 µl pre-warmed SOC and incubated while shaking at 37°C for 45 min. We targeted ∼100-500 colonies per variant to ensure that sufficient 16N barcodes would be produced in the final library. We estimated transformant counts by generating spot plates from the transformed culture. Specifically, we spotted 2 uL dilutions (ranging from 10^-1^ - 10^-3^ initial concentration) in triplicate on selection plates which we incubated overnight at 37°C. Simultaneously, we added growth media (LB + 100 µg/mL carbenicillin) to 50 µL, 100 µL and 500 µL aliquots of the transformed culture. These tubes were incubated overnight while shaking at 37 °C. After overnight growth, the transformation efficiency was calculated based on the spot plate colony counts. This efficiency measurement was used to determine that the 100 µL sample would provide approximately ∼100-500 16N barcodes per variant. The 100 µL sample was miniprepped, yielding the full recombination-ready donor plasmid library. A glycerol stock (25% v/v sterile glycerol, -80°C storage) of the culture was created simultaneously.

### Sequencing of PROD-ATAC donor plasmid libraries prior to recombination

We sequenced the random 16N barcodes and the constant 12-nt barcodes after insertion into the plasmid attB-containing backbone with a 1 × 150-bp single-end sequencing run. We filtered for correctly structured amplicons by grepping for the four following constant sequences which flanked the 12-nt and 16N barcodes: ‘AAATCCAAGC’, ‘CCAGAGCATG’, ‘CAAGGTGGTT’ and ‘ATACTGATTC’. We first generated a list of 12-nt constant – 16N random barcode pairs. Next, we restricted the list of pairs to those that were seen more than once. Finally, we required that, for a given random barcode, more than 95% of the reads matched the same constant 12-nt barcode. Should a random barcode frequently map to more than one hardcoded barcode, we would not assign said barcode to a particular perturbation.

### Library recombination and sequencing

To create the full vTR library, 293T-LP cells were thawed and passaged twice before recombination. Three replicate wells of a six-well plate were seeded at 400,000 cells per well 18 hrs prior to transfection. The cultures were then recombined with the plasmid attB-containing library. All three of the six wells were transfected with 100 ng of pCAG-NLS-HA-Bxb1 (Addgene, 51271) and 1.5 µg of the donor vTR library. Transfection was carried out using Lipofectamine 3000 with serum-free Opti-Mem diluent. Liposomes were added to cells at approximately 50% confluence and incubated for 18 hrs. After incubation, liposome-containing media was aspirated and replaced with fresh complete media containing puromycin at 0.8 µg/mL to select for recombined cells. All recombined 293T-LP cell populations were hereafter maintained in puromycin-containing media. After 3 d of puromycin selection, all cultures were combined and scaled up to a single T75 culture seeded with 3.0 × 10^6^ cells. These cells were then maintained in selection-conditions with periodic passages that prevented cells from reaching >90% confluency. After 11 d of selection and culture maintenance, we froze down several vials with 2 million cells each. The cell population gDNA was isolated using a PureLink Genomic DNA Mini Kit for downstream analyses (proliferation assays, variant representation).

The condensed 15-member vTR library was recombined and sequenced in the exact same manner, merely using a donor plasmid library with the target 15 (rather than full 110) variant sequences.

### Engineering arrayed 293T-LP and K562 vTR cell lines for clonal assays

We generated individual donor plasmids encoding five different variants chosen from the initial library. For each variant, we performed the identical PCR, Golden Gate Assembly, and transformation described in the library assembly steps. We purified plasmids from single colonies picked for each variant and verified sequence correctness via full plasmid sequencing through Plasmidsaurus. We then recombined the donor plasmids into the 293T-LP cell lines to generate five clonal cell lines each singly expressing the variant (EV, LMP1, BGLF4, Tax (HTLV-1), and Zta). Again, these cell lines were generated in the same strategy as described for library creation. Successful recombination was verified by puromycin resistance and PCR of the inserted variant sequences from harvested gDNA.

Separately, we generated two K562 clonal cell lines that expressed either EV control or E4orf6/7 (HAdvB). To create these cell lines, we used lentiviral transgene delivery rather than the previously described attB donor constructs (as the K562 cell line lacked our specific attP-landing site). First, we thawed 293T cells and passaged them twice ensuring they never achieved >90% confluency. Two wells of 6-well plates per variant (EV and E4orf6/7) were seeded with 450,000 cells 24 hrs prior to transfection with lentiviral assembly plasmids. We then transfected each well with 1.3 µg psPAX2 (Addgene, 12260), 200 ng pCMV-VSV-G (Addgene, 8454), and 1.4 µg of donor plasmid (pCDH-Tet-Puro) that encoded either the EV or E4orf6/7 cassette. Transfection was performed using Lipofectamine 3000 with Opti-Mem media. Liposome-containing media was aspirated 18 hrs post-transfection and replaced with fresh pre-warmed media. The lentivirus-containing supernatant from each well was harvested 72 hrs after media replacement. The supernatants were then spun down at 500xg for 5 min and sterilely filtered (0.45 µm pore size). Approximate lentiviral titers were subsequently determined by applying dilutions of the filtered virus-containing supernatant to 293T cells and quantifying resistant cells.

K562 cells were then transduced with the prepared lentiviral populations. Wells were seeded at a density of 100,000 cells / well 18 hrs prior to transduction. The EV and E4orf6/7-packaged lentivirus populations were next applied to the K562 cells. For each transduction, we targeted a sub-0.1 multiplicity of infection (MOI) to ensure cells were at most singly infected (based on Poisson distribution). After 24 hr, we replaced the cell media with fresh complete RPMI supplemented with 2 µg/mL puromycin to select for transduced K562 cells. All transduced K562 population were hereafter maintained in puromycin-containing media. The transduced K562 wells were passaged routinely and maintained at a density between 200,000-900,000 cells/mL. After 12 days in selection conditions, several samples of 2 million cells each were frozen to be thawed later for bulk ATAC experiments.

### Pooled library proliferation screen

To test whether expression of any of the variants had a proliferative effect and, therefore, altered the library distribution, we performed a pooled screen of the 110-member 293T-LP cell library. We seeded six replicate wells of the library. Three wells were induced with doxycycline (2 µg/mL), whereas the other three were treated with an equal volume of water. Replicates were passaged 48 hrs post-induction. At 96 hrs post-induction, 500,000 cells from each sample (triplicate water and doxycycline-treated conditions) were used for gDNA isolation and barcode sequencing to quantify variant distribution. The gDNA NGS and barcode filtering were performed as described above.

### Single-cell ATAC and dial-out library preparation

The full-scale (110-member) PROD-ATAC library was generated as detailed previously^23^. Briefly, 293T-LP cells encoding the 110-member library were seeded in a T-25 flask and allowed to attach overnight before being induced with doxycycline (2 µg/mL) for 96 hr with one split at 48 hr. For this work, all sequencing was performed using 10x Genomics v2 Chemistry. Nuclei were prepared according to 10x Genomics’ Nuclear Isolation for Single Cell ATAC Sequencing protocol with the following changes. A 10% solution IGEPAL CA-630 solution was prepared from a 100% solution (Sigma-Aldrich, i8896) to be used for lysis buffer preparation. Following a 4 min lysis period, cells were resuspended in 1x nuclei buffer. Triplicates were counted before proceeding with PROD-ATAC sequencing as previously published. In all cases, 18,000 nuclei were targeted for capture per 10x chip lane. The full-scale PROD-ATAC vTR screen used one full 10x chip, which includes 8 lanes (for a target of ∼140,000 total cells from the same population). Single-cell ATAC library fragment size distribution for each 10x chip lane was analyzed by TapeStation before preparing enriched dial-out libraries for variant genotyping as previously detailed.

For the PROD-ATAC drug perturbation assays, the following adaptations were made to the above workflow. 293T-LP cells encoding the 15-member condensed library were seeded into three separate wells of a six-well plate and allowed to attach overnight. After 18 hrs, these cells were induced with doxycycline (2 µg/mL). Simultaneously, the three wells were separately treated with JQ1 (50 nM), TAK-733 (100 nM), or a vehicle control (0.1% DMSO). The three cultures were treated in the same manner as detailed above until 10x chip loading. At this stage, each condition was added to a separate 10x chip lane, with a target of 18,000 cells / lane. The scATAC and dialout libraries were then created by the same procedure as described for the full vTR library.

### Single-cell ATAC data processing

PROD-ATAC data analysis for all single-cell ATAC libraries and associated genotyping dial-out libraries was performed as previously described^23^. Briefly, single-cell sequencing datafiles were converted to FASTQ format using cellranger-atac mkfastq (10x Genomics, version 2.1.0) and aligned to the hg38 reference genome via cellranger-atac count. ArchR (version 1.0.3) was used for calling vTR marker peaks and downstream analysis using the same filter settings and quality cutoffs as described previously. Fragment files generated via cellranger-atac count (along with associated cell-assignment file outputs) were used to build Arrow files in ArchR.

ArchR (version 1.0.3)^30^ was used for calling vTR marker peaks and performing downstream analysis as described in depth previously^23^. Briefly, all nuclei used for PROD-ATAC data analysis were first filtered for those with TSS enrichment scores ≥6 and ≥15,000 fragments. We then generated pseudobulk samples representing each vTR genotype and created in silico replicates to perform statistical tests to identify differentially accessible peaks. The addGroupCoverages command (minCells = 20, maxCells = 500, minReplicates = 2, maxReplicates = 5) merged cells of a known genotype. The resultant coverage file was queried to identify peaks via addReproduciblePeakSet. MACS3 was used with default parameters for peak calling^33^. Patterns of motif enrichment within these differentially accessible peaks were identified by calling peakAnnoEnrichment with default parameters.

### Linking vTR genotypes to ATAC datasets via dial-out library sequencing

Dial-out enrichment libraries used to call perturbation identities in captured nuclei were sequenced as in our previous oncofusion PROD-ATAC study^23^. Briefly, the random 16N barcodes were sequenced and mapped to the 12-nt variant-specific barcodes. This therefore linked the 16N barcodes, ATAC libraries, and associated vTR sequences. Random barcodes were again identified by grepping for four constant sequences (‘AAATCCAAGC’, ‘CCAGAGCATG’, ‘CAAGGTGGTT’ and ‘ATACTGATTC’) to isolate correctly arranged amplicons. Random barcodes were then isolated alongside their corresponding cell barcodes (CBCs) from the 10x library construction process. We filtered on 10x CBCs that were seen at least three times and retained pairs of 10x CBC-random barcode for which ≥90% of the reads were identical. Random barcodes from the dial-out library were matched to random barcodes identified from sequencing the library from gDNA previously. The remainder of cell barcodes that were unidentified (ie. did not find matches in the known 16N barcode list) were excluded from downstream analysis.

### Bulk RNA-seq sample preparation and sequencing

We generated bulk RNA-seq datasets from nine clonal cell lines (for eight vTRs + EV control). The 293T-LP cells were treated in the same manner as described for the bulk ATAC sequencing assays. Each variant was seeded in two wells of 6-well plates to generate biological duplicates. After 96 hrs of doxycycline induction (as for all ATAC samples), 200,000 cells from each condition were harvested and lysed in 50 µL Zymo DNA/RNA Shield™. As the samples were highly viscous, each lysed sample was vigorously vortexed for 15-25 seconds to shear chromosomes and facilitate downstream sample preparation. Each sample (18x total for biological duplicates) was then shipped to Plasmidsaurus for 3’ RNA-seq library generation.

RNA-seq libraries were prepared and sequenced by Plasmidsaurus using their standard 3′ end counting workflow. Briefly, total RNA was isolated from each lysed biological replicate and poly-A selection with oligo-dT primers yielded mRNA-specific libraries. Reverse transcription and second-strand synthesis generated double-stranded cDNA. The cDNA libraries were tagmented and finally amplified to incorporate unique sample indices and Illumina adapter sequences for NGS. The Plasmidsaurus workflow also inserts unique molecular identifiers (UMIs) during cDNA synthesis to enable read de-duplication. Each vTR RNA sample targeted approximately 10 million de-duplicated reads per sample. Sequencing reads (derived from the 3′ transcript ends) were subsequently analyzed to generate differential gene expression counts.

### Bulk ATAC sample analyses

Bulk ATAC analyses were performed as previously described^23^. Paired-end reads were first merged with NGmerge in adapter removal mode (-a -e 20). Forward and reverse reads were separately aligned to the same reference genome used for our single-cell sequencing analysis (refdata-cellranger-arc-GRCh38-2020-A-2.0.0) using the same BWA framework for SAM to BAM file generation. BAM files were filtered to eliminate reads mitochondrial-aligned reads (SAMtools idxstats bam_file.bam | cut -f 1 | grep -v NC_012920.1 | xargs samtools view - b bam_file.bam > bam_file_no_mito.bam). We next calculated Pearson correlations for all pairwise comparisons. To do this, we merged the two BAM files for each vTR (SAMtools merge) and created BigWig tracks for each combined sample (bamCoverage–normalizeUsing CPM). We then calculated the average score either over all genome bins or over specifically defined regions with multiBigwigSummary. Outliers were excluded when calculating correlations (plotCorrelation–removeOutliers -c pearson). In Fig. 6a, dynamic sites were defined as those that were differentially accessible in pseudobulk comparisons of BGLF4, Zta, Tax (HTLV-1), and LMP1 to the EV control.

### vTR AlphaFold3 structural predictions

Predicted three-dimensional vTR structures were generated through the publicly accessible AlphaFold3 interface using default parameters^69^. Briefly, amino acid sequences for the selected vTRs were retrieved from UniProt and input into the AlphaFold3 prediction pipeline. We produced five separate structural models per vTR sequence. We used Predicted Local Distance Difference Test (pLDDT) scores and Predicted Aligned Error (PAE) heatmaps to determine modeling fidelity and to identify protein regions with sufficient confidence to be reliably interpreted. The highest-confidence monomer structure generated for each vTR was selected for visualization and structural similarity comparisons.

### Mammalian-one-hybrid (M1H) assays

We measured transcriptional activity of the Tax (HTLV-1/2/3) homologs using M1H assays in 293T cells. For these assays, vTRs were cloned into the DB-pEZY3 vector via Gateway cloning such that the N-terminus of each variant was fused to the Gal4 DNA-binding domain^7^. M1H assays were performed using pGL4.23 luciferase reporter constructs. The pGL4.23 constructs encoded a moderate promoter, 4x upstream activating sequences (UAS), and the Syn2B10 enhancer sequence.

293T cells were plated in 96-well white opaque plates (Corning) at a density of 10,000 cells/well and incubated for 24h. Cells were then transfected with 80 ng of the vTR DB-pEZY3 construct, 20 ng 4xUAS construct, and 10 ng of renilla luciferase plasmid as a transfection normalization control. An empty DB-pEZY3 plasmid and 4xUAS construct plasmid were co-transfected to serve as a negative control. Cells were incubated for 48hr after transfection, followed by measurement of both firefly and renilla luciferase activity using the Dual-Glo Luciferase Assay System (Promega). Non-transfected cells were used to subtract background for firefly/renilla luciferase activities. Firefly luciferase activity was then normalized with renilla luciferase activity in each well.

### Genome annotations

All analyses for 293T-LP and K562 datasets were performed using the reference hg38 genome. Gene annotations for RNA-seq analyses were derived from “TxDb.Hsapiens.UCSC.hg38.knownGene”.

## Code availability

Scripts relevant for downstream PROD-ATAC data processing and figure generation used in both this and other PROD-ATAC associated publications are deposited in GitHub (https://github.com/mfrenkel16/OncofusionPRODATAC/).

## References

1. Liang, G. & Bushman, F. D. The human virome: assembly, composition and host interactions. Nat. Rev. Microbiol. 19, 514–527 (2021).

2. Harvey, E. & Holmes, E. C. Diversity and evolution of the animal virome. Nat. Rev. Microbiol. 20, 321–334 (2022).

3. Dion, M. B., Oechslin, F. & Moineau, S. Phage diversity, genomics and phylogeny. Nat. Rev. Microbiol. 18, 125–138 (2020).

4. Prangishvili, D., Forterre, P. & Garrett, R. A. Viruses of the Archaea: a unifying view. Nat. Rev. Microbiol. 4, 837–848 (2006).

5. Liu, X. et al. Human Virus Transcriptional Regulators. Cell 182, 24–37 (2020).

6. Arvey, A. et al. An atlas of the Epstein-Barr virus transcriptome and epigenome reveals host-virus regulatory interactions. Cell Host Microbe 12, 233–245 (2012).

7. Rottenberg, J. T. et al. Viral transcriptional regulators extensively rewire host pathways through diverse mechanisms. 2025.12.01.691387 Preprint at 10.64898/2025.12.01.691387 (2025).

8. Tsai, K. & Cullen, B. R. Epigenetic and epitranscriptomic regulation of viral replication. Nat. Rev. Microbiol. 18, 559–570 (2020).

9. Uppal, T., Banerjee, S., Sun, Z., Verma, S. C. & Robertson, E. S. KSHV LANA—The Master Regulator of KSHV Latency. Viruses 6, 4961–4998 (2014).

10. Matsuoka, M. & Jeang, K.-T. Human T-cell leukaemia virus type 1 (HTLV-1) infectivity and cellular transformation. Nat. Rev. Cancer 7, 270–280 (2007).

11. Davis, Z. H. et al. Global mapping of herpesvirus-host protein complexes reveals a transcription strategy for late genes. Mol. Cell 57, 349–360 (2015).

12. Kosmopoulos, J. C. & Anantharaman, K. Viral Dark Matter: Illuminating Protein Function, Ecology, and Biotechnological Promises. Biochemistry 64, 4609–4627 (2025).

13. Kim, K.-D. & Lieberman, P. M. Viral remodeling of the 4D nucleome. Exp. Mol. Med. 56, 799–808 (2024).

14. Yu, Y., Mai, Y., Zheng, Y. & Shi, L. Assessing and mitigating batch effects in large-scale omics studies. Genome Biol. 25, 254 (2024).

15. Leek, J. T. et al. Tackling the widespread and critical impact of batch effects in high-throughput data. Nat. Rev. Genet. 11, 733–739 (2010).

16. Morris, J. A., Sun, J. S. & Sanjana, N. E. Next-generation forward genetic screens: uniting high-throughput perturbations with single-cell analysis. Trends Genet. TIG 40, 118–133 (2024).

17. High-content CRISPR screening. Nat. Rev. Methods Primer 2, 9 (2022).

18. Dixit, A. et al. Perturb-seq: Dissecting molecular circuits with scalable single cell RNA profiling of pooled genetic screens. Cell 167, 1853–1866.e17 (2016).

19. Pierce, S. E., Granja, J. M. & Greenleaf, W. J. High-throughput single-cell chromatin accessibility CRISPR screens enable unbiased identification of regulatory networks in cancer. Nat. Commun. 12, 2969 (2021).

20. Ursu, O. et al. Massively parallel phenotyping of coding variants in cancer with Perturb-seq. Nat. Biotechnol. 40, 896–905 (2022).

21. Parekh, U. et al. Mapping Cellular Reprogramming via Pooled Overexpression Screens with Paired Fitness and Single-Cell RNA-Sequencing Readout. Cell Syst. 7, 548–555.e8 (2018).

22. Xie, S., Cooley, A., Armendariz, D., Zhou, P. & Hon, G. C. Frequent sgRNA-barcode recombination in single-cell perturbation assays. PLoS ONE 13, e0198635 (2018).

23. Frenkel, M., Corban, J. E., Hujoel, M. L. A., Morris, Z. & Raman, S. Large-scale discovery of chromatin dysregulation induced by oncofusions and other protein-coding variants. Nat. Biotechnol. 43, 996–1010 (2025).

24. Krump, N. A. & You, J. Molecular mechanisms of viral oncogenesis in humans. Nat. Rev. Microbiol. 16, 684–698 (2018).

25. Law, G. L., Korth, M. J., Benecke, A. G. & Katze, M. G. Systems virology: host-directed approaches to viral pathogenesis and drug targeting. Nat. Rev. Microbiol. 11, 455–466 (2013).

26. Chen, I. P. & Ott, M. Viral Hijacking of BET Proteins. Viruses 14, 2274 (2022).

27. DuShane, J. K. & Maginnis, M. S. Human DNA Virus Exploitation of the MAPK-ERK Cascade. Int. J. Mol. Sci. 20, 3427 (2019).

28. Hill, A. J. et al. On the design of CRISPR-based single cell molecular screens. Nat. Methods 15, 271–274 (2018).

29. Ramasubramanyan, S. et al. Epstein–Barr virus transcription factor Zta acts through distal regulatory elements to directly control cellular gene expression. Nucleic Acids Res. 43, 3563–3577 (2015).

30. Granja, J. M. et al. ArchR is a scalable software package for integrative single-cell chromatin accessibility analysis. Nat. Genet. 53, 403–411 (2021).

31. Hao, Y. et al. Integrated analysis of multimodal single-cell data. Cell 184, 3573–3587.e29 (2021).

32. Jost, M. et al. Titrating gene expression using libraries of systematically attenuated CRISPR guide RNAs. Nat. Biotechnol. 38, 355–364 (2020).

33. Zhang, Y. et al. Model-based Analysis of ChIP-Seq (MACS). Genome Biol. 9, R137 (2008).

34. Pantry, S. N. & Medveczky, P. G. Epigenetic regulation of Kaposi’s sarcoma-associated herpesvirus replication. Semin. Cancer Biol. 19, 153–157 (2009).

35. Weigel, C. et al. Epigenetic activation of EBV BGLF4 determines antiviral-based regimen response in EBV+CNS lymphoproliferative disease. Blood Neoplasia 3, 100200 (2026).

36. Martin, K. A., Lupey, L. N. & Tempera, I. Epstein-Barr Virus Oncoprotein LMP1 Mediates Epigenetic Changes in Host Gene Expression through PARP1. J. Virol. 90, 8520–8530 (2016).

37. Romanelli, M. G. et al. Highlights on distinctive structural and functional properties of HTLV Tax proteins. Front. Microbiol. 4, (2013).

38. Damania, B. & Jacobs, S. R. The Viral Interferon Regulatory Factors of KSHV: Immunosuppressors or Oncogenes? Front. Immunol. 2, (2011).

39. Schaley, J. E., Polonskaia, M. & Hearing, P. The Adenovirus E4-6/7 Protein Directs Nuclear Localization of E2F-4 via an Arginine-Rich Motif. J. Virol. 79, 2301–2308 (2005).

40. Dexheimer, P. J. et al. Systematic investigation reveals extensive Epstein-Barr virus transcriptional regulation of the human genome. 2025.07.18.665561 Preprint at 10.1101/2025.07.18.665561 (2025).

41. Chang, L.-S. et al. Epstein-Barr Virus BGLF4 Kinase Downregulates NF-κB Transactivation through Phosphorylation of Coactivator UXT. J. Virol. 86, 12176–12186 (2012).

42. Zhao, T. The Role of HBZ in HTLV-1-Induced Oncogenesis. Viruses 8, 34 (2016).

43. Terhune, S. S. et al. Human cytomegalovirus UL29/28 protein interacts with components of the NuRD complex which promote accumulation of immediate-early RNA. PLoS Pathog. 6, e1000965 (2010).

44. Hübner, M. R., Eckersley-Maslin, M. A. & Spector, D. L. Chromatin organization and transcriptional regulation. Curr. Opin. Genet. Dev. 23, 89–95 (2013).

45. Di, C. et al. RTA and LANA Competitively Regulate let-7a/RBPJ Signal to Control KSHV Replication. Front. Microbiol. 12, 804215 (2022).

46. Luftig, M. et al. Epstein–Barr virus latent membrane protein 1 activation of NF-κB through IRAK1 and TRAF6. Proc. Natl. Acad. Sci. 100, 15595–15600 (2003).

47. Herbein, G., Gras, G., Khan, K. A. & Abbas, W. Macrophage signaling in HIV-1 infection. Retrovirology 7, 34 (2010).

48. Barrado-Gil, L., Galindo, I., Martínez-Alonso, D., Viedma, S. & Alonso, C. The ubiquitin-proteasome system is required for African swine fever replication. PLOS ONE 12, e0189741 (2017).

49. Barrado-Gil, L. et al. African Swine Fever Virus Ubiquitin-Conjugating Enzyme Is an Immunomodulator Targeting NF-κB Activation. Viruses 13, 1160 (2021).

50. El-Guindy, A., Lopez-Giraldez, F., Delecluse, H.-J., McKenzie, J. & Miller, G. A Locus Encompassing the Epstein-Barr Virus bglf4 Kinase Regulates Expression of Genes Encoding Viral Structural Proteins. PLoS Pathog. 10, e1004307 (2014).

51. Karmakar, S. & Reilly, K. M. The role of the immune system in neurofibromatosis type 1-associated nervous system tumors. CNS Oncol. 6, 45–60 (2017).

52. Lin, Y.-C. et al. Genome dynamics of the human embryonic kidney 293 lineage in response to cell biology manipulations. Nat. Commun. 5, 4767 (2014).

53. Yamano, S. et al. Induction of Transformation and p53-Dependent Apoptosis by Adenovirus Type 5 E4orf6/7 cDNA. J. Virol. 73, 10095–10103 (1999).

54. Zhou, B. et al. Comprehensive, integrated, and phased whole-genome analysis of the primary ENCODE cell line K562. Genome Res. 29, 472–484 (2019).

55. Ludwig, C. H. et al. High-throughput discovery and characterization of viral transcriptional effectors in human cells. Cell Syst. 14, 482–500.e8 (2023).

56. Liberzon, A. et al. The Molecular Signatures Database (MSigDB) hallmark gene set collection. Cell Syst. 1, 417–425 (2015).

57. Wang, J., Zhou, J.-Y., Kho, D., Reiners, J. J. & Wu, G. S. Role for DUSP1 (dual-specificity protein phosphatase 1) in the regulation of autophagy. Autophagy 12, 1791–1803 (2016).

58. Boiko, A. D. et al. A systematic search for downstream mediators of tumor suppressor function of p53 reveals a major role of BTG2 in suppression of Ras-induced transformation. Genes Dev. 20, 236–252 (2006).

59. Hao, Y. et al. Integrated analysis of multimodal single-cell data. Cell 184, 3573–3587.e29 (2021).

60. Dong, Q. et al. Discovery of TAK-733, a potent and selective MEK allosteric site inhibitor for the treatment of cancer. Bioorg. Med. Chem. Lett. 21, 1315–1319 (2011).

61. Li, Z., Guo, J., Wu, Y. & Zhou, Q. The BET bromodomain inhibitor JQ1 activates HIV latency through antagonizing Brd4 inhibition of Tat-transactivation. Nucleic Acids Res. 41, 277–287 (2013).

62. Teo, A. Y. Y., Squair, J. W., Courtine, G. & Skinnider, M. A. Best practices for differential accessibility analysis in single-cell epigenomics. Nat. Commun. 15, 8805 (2024).

63. Filippakopoulos, P. et al. Selective inhibition of BET bromodomains. Nature 468, 1067–1073 (2010).

64. Che, X., Zerboni, L., Sommer, M. H. & Arvin, A. M. Varicella-Zoster Virus Open Reading Frame 10 Is a Virulence Determinant in Skin Cells but Not in T Cells In Vivo. J. Virol. 80, 3238–3248 (2006).

65. Kinnunen, M. et al. The Impact of ETV6-NTRK3 Oncogenic Gene Fusions on Molecular and Signaling Pathway Alterations. Cancers 15, 4246 (2023).

66. Qiu, T. et al. CCDC6-RET fusion protein regulates Ras/MAPK signaling through the fusion-GRB2-SHC1 signal niche. Proc. Natl. Acad. Sci. U. S. A. 121, e2322359121.

67. Leek, J. T. et al. Tackling the widespread and critical impact of batch effects in high-throughput data. Nat. Rev. Genet. 11, 733–739 (2010).

68. Citu, C., Chang, L., Manuel, A. M., Enduru, N. & Zhao, Z. Identification and catalog of viral transcriptional regulators in human diseases. iScience 28, 112081 (2025).

69. Abramson, J. et al. Accurate structure prediction of biomolecular interactions with AlphaFold 3. Nature 630, 493–500 (2024).

