## Supplemental figures for "Systematic mapping of chromatin dysregulation driven by viral transcriptional regulators at scale"

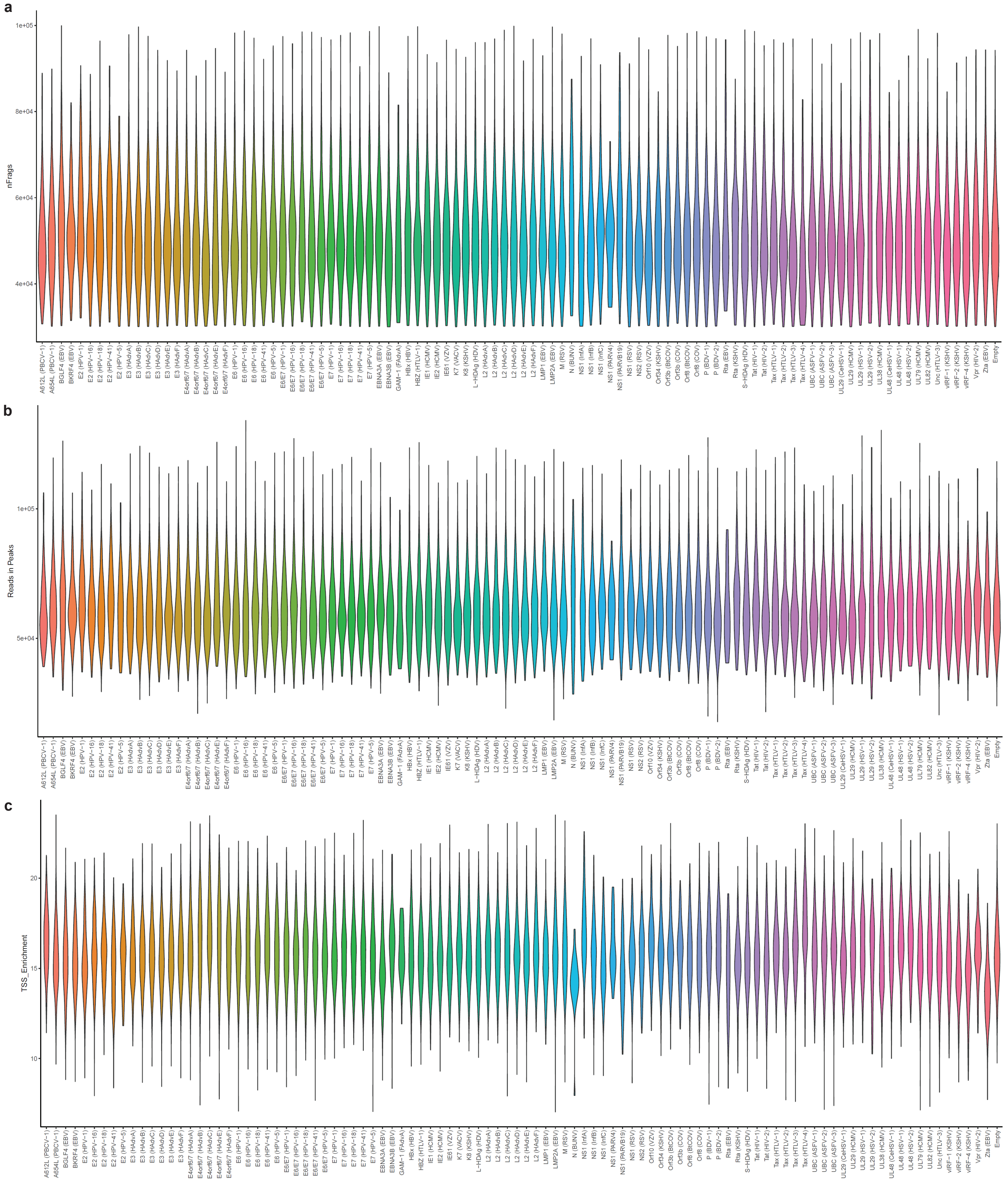


**Supplementary Figure 1: Nuclei and ATAC libraries corresponding to vTR variants are high quality and comparable in coverage. (a)** Distribution of fragments captured for nuclei associated with each variant. **(b)** Distribution of reads in peak regions for nuclei associated with each variant. **(c)** Distribution of TSS (transcription start site) enrichment scores for nuclei associated with each variant.

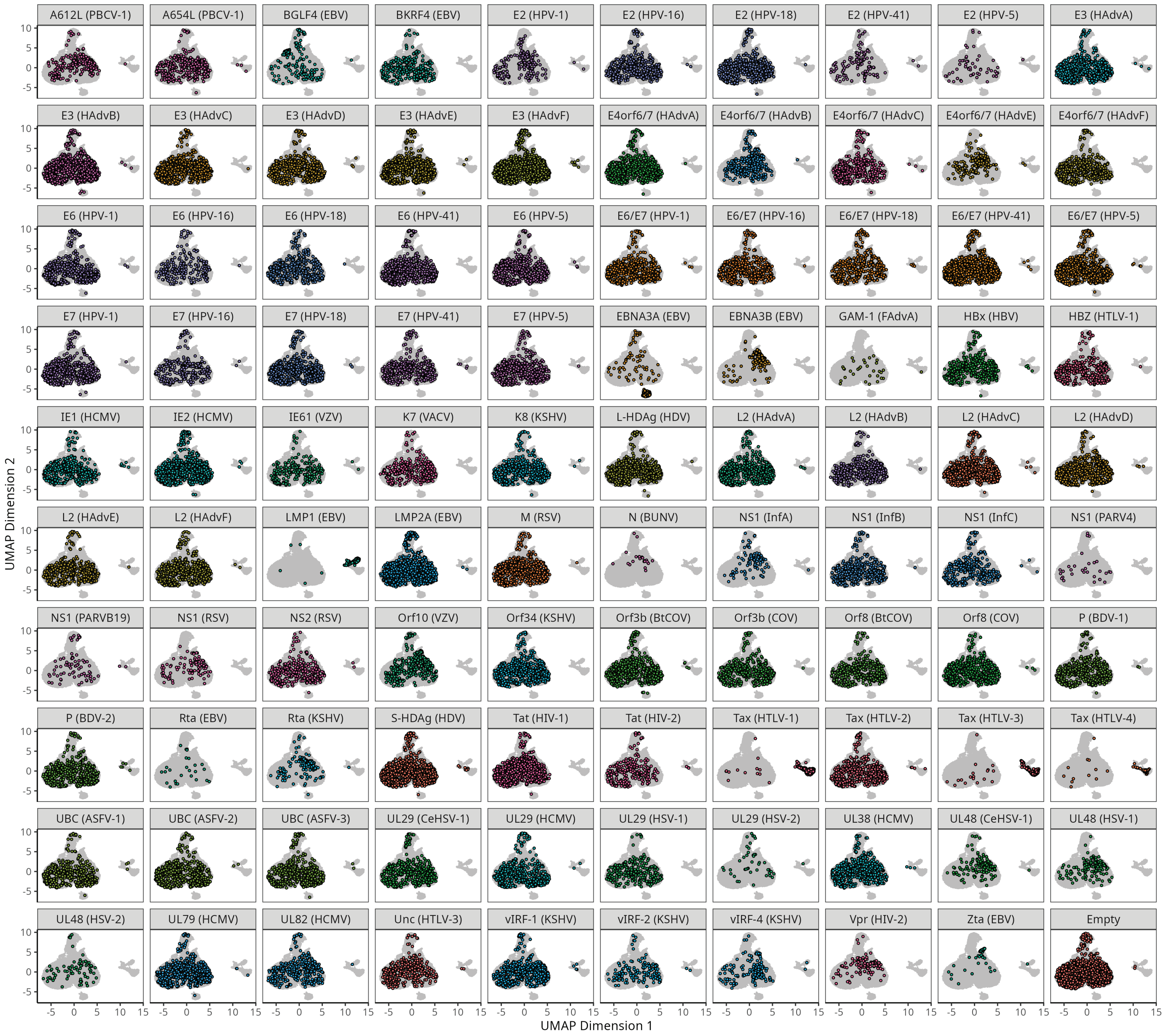


**Supplementary Figure 2: Chromatin accessibility-driven distributions of variant-genotyped nuclei in UMAP space.** All 43,798 genotyped nuclei in the full-scale vTR PROD-ATAC screen are displayed in UMAP space post-LSI (latent semantic indexing). Each subpanel shows the nuclei genotyped to a given vTR variant highlighted against the rest of the combined assayed nuclei (shown in grey).

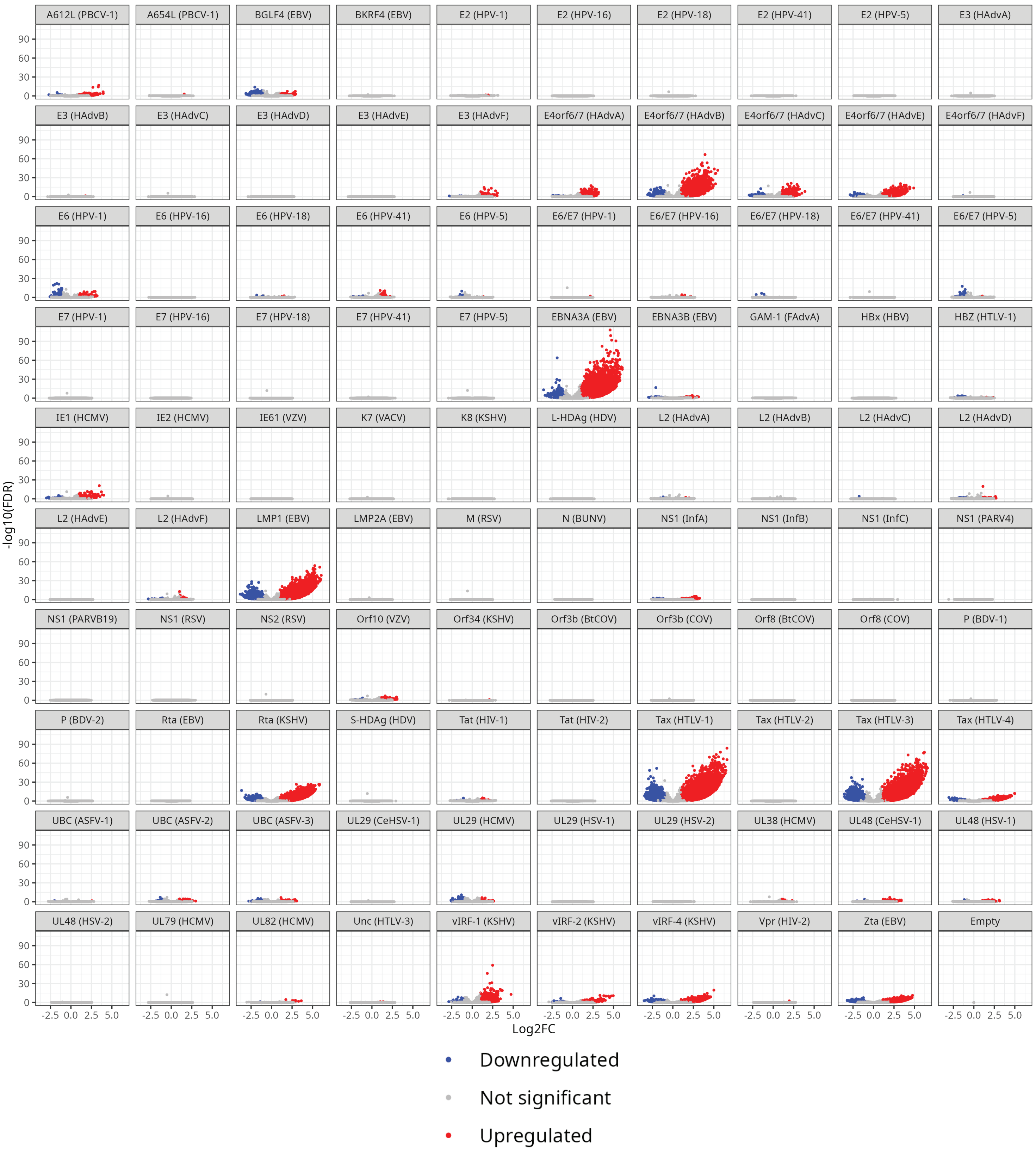


**Supplementary Figure 3: Chromatin dysregulation measurements of each vTR in full-scale PROD-ATAC screen.** Volcano plots for all vTR variants (including all variants which produced no significantly dysregulated peaks) comparing pseudobulk replicates to EV control, with −log(FDR) versus $\log_2$FC displayed in each subplot for called peaks. Increased accessibility peaks (FDR $\leq$ 0.1 and $\log_2$FC $\geq$ 1) are colored red, and decreased accessibility peaks (FDR $\leq$ 0.1 and $\log_2$FC $\leq$ −1) are colored blue.

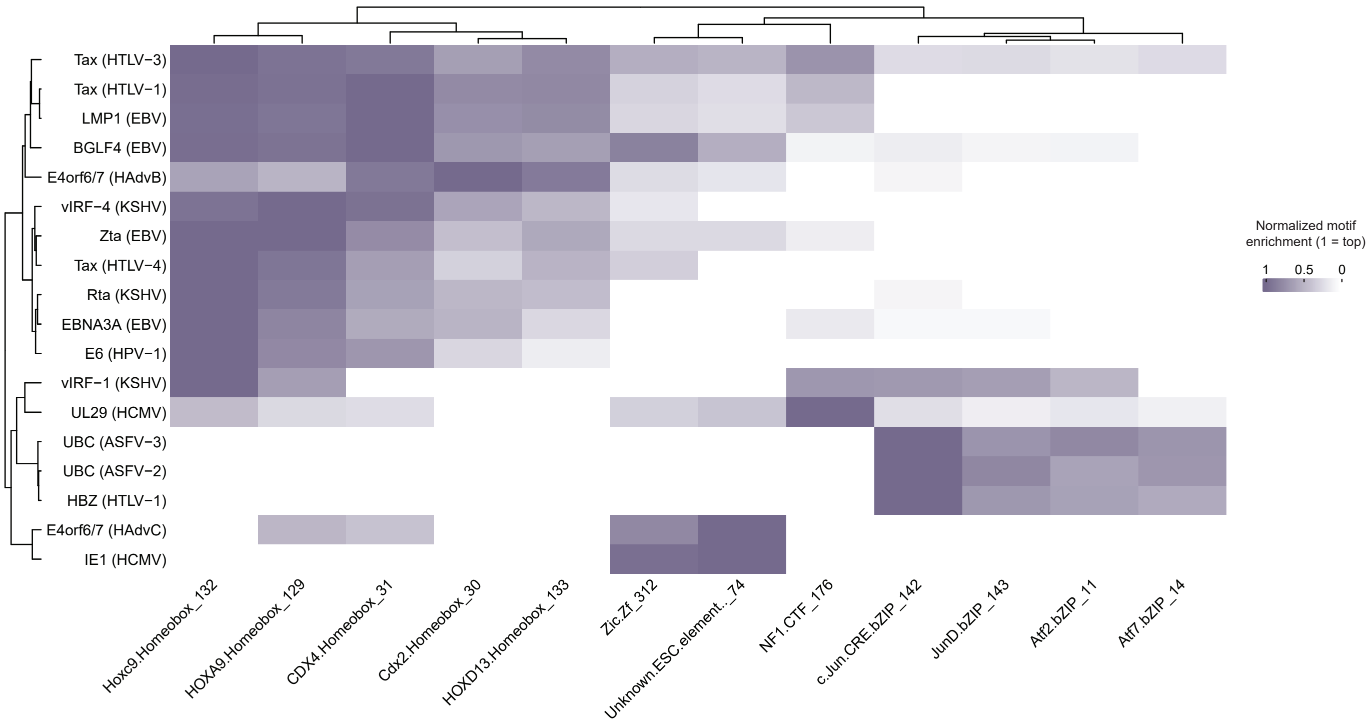


**Supplementary Figure 4: Chromatin regions closed through activity of diverse vTRs converge on similar hTF motif enrichments**. Motif enrichment for vTRs with significantly enriched DNA motifs in decreased accessibility peaks. Only variants with at least one substantially enriched motif (−log(*P*) > 5)) and only motifs enriched in at least one set of vTR pseudobulk replicates are shown. TF motifs and vTRs are each clustered by Euclidean distance.
